# DNA methylome responses to biotic and abiotic stress in *Arabidopsis thaliana*: A multi-study analysis

**DOI:** 10.1101/2025.10.20.682861

**Authors:** R Behl, JJ Gallo-Franco, RR Hazarika, Z Zhang, F Wilming, JP Schnitzler, C Lindermayr, F Johannes

**Author notes:** These authors contributed equally to this work (R Behl, JJ Gallo-Franco). Email addresses.

## Abstract

Plants have evolved complex physiological and morphological strategies to cope with biotic and abiotic stress. At the molecular level, these responses are partly mediated by epigenetic mechanisms such as DNA methylation. In *Arabidopsis thaliana*, numerous studies have examined stress-induced changes in DNA methylation, aiming to elucidate their roles in stress acclimation. However, comparing these studies remains difficult due to variations in analytical methods and focus. To address this, we conducted an integrative analysis of 13 studies (16 datasets) using a unified bioinformatic pipeline to identify common and stress-specific methylation responses. Our results show that the number and distribution of differentially methylated regions (DMRs) vary by stress type and methylation context CG, CHG, and CHH. CG DMRs were primarily located in gene bodies, while CHG and CHH DMRs were enriched in transposable elements (TEs). Gene Ontology analysis revealed consistently enriched stress-related terms across conditions, indicating shared epigenetic regulation. As TEs have been proposed to act as stress sensors and regulators of nearby gene expression, we examined gene-proximal TEs in more detail. We found significant enrichment of stress-responsive methylation changes in *LINE/L1, RathE1_ons,* and *DNA/HAT* elements. While stress-induced methylation is often considered transient, many affected CG DMRs overlapped with loci known to accumulate stable epimutations over generations, suggesting that stress may contribute to long-term heritable epigenetic variation. These findings enhance our understanding of epigenetic regulation in plant stress responses, highlighting both shared and stress-specific methylation dynamics.

**Significance statement:** We combined 16 *Arabidopsis thaliana* Whole-Genome Bisulfite Sequencing datasets to reveal common and stress-specific DNA methylation responses. Although global methylation levels remained stable, specific DMRs varied according to the type of stress and genomics context. Gene Ontology analysis revealed shared stress pathways. Gene-proximal transposable elements (TEs) (e.g. LINE/L1, RathE1_ons, DNA/HAT) were enriched for stress-responsive methylation changes. Notably, a subset of stress-induced CG DMRs overlap with epimutation-prone loci, suggesting an environmental route toward stable, heritable epigenetic variation.

## Introduction

Climatic changes are increasing the frequency and intensity of environmental stresses (Schillerberg and Tian, 2024, Gebrechorkos et al., 2025). As sessile organisms, plants must rapidly sense and adapt to a wide variety of abiotic and biotic stressors, ranging from drought, salinity, and extreme temperatures to pathogen attack (Jiang et al., 2025). A key mechanism underlying these adaptive responses is DNA methylation, an epigenetic modification that regulates gene expression and TE activity (Law and Jacobsen, 2010). In plants, methylation occurs in CG, CHG, and CHH sequence contexts (where H = A, T, C) and is catalyzed by distinct sets of DNA methyltransferases (e.g., MET1, CMT2/3, DRM2), while active demethylation is carried out by enzymes such as ROS1 and DML2/3 (Finnegan and Kovac, 2000, Finnegan et al., 2000, Dalakouras et al., 2012, Zhang et al., 2018, Sun et al., 2022).

DNA methylation plays diverse roles depending on its genomic context. In gene promoters, it typically represses transcription by blocking transcription factor access, while gene body methylation (gbM) - usually in the CG context - is associated with moderately expressed housekeeping genes and may stabilize transcription (Muyle et al., 2022). Methylation within TEs is especially important for silencing their activity and maintaining genome integrity. Emerging evidence also suggests that gene-proximal TEs have been co-opted during evolution and domestication as environmentally responsive regulatory elements (Dubin et al., 2018, Roquis et al., 2021, Ito, 2022). These elements may act as tunable modulators of gene expression, with stress-induced differential methylation serving as a mechanism for fine-tuning transcriptional responses to environmental cues (Dubin et al., 2015).

A growing body of evidence supports a regulatory role of DNA methylation in stress acclimation (Kumar and Mohapatra, 2021, Zhang et al., 2018, Bräutigam and Cronk, 2018). For instance, in *Arabidopsis thaliana*, salt stress leads to demethylation of the AtMYB74 promoter, resulting in its transcriptional activation - a response likely essential for stress tolerance (Xu et al., 2015, Bartels et al., 2018). In the context of biotic stress, pathogen exposure can trigger dynamic shifts in CHH methylation and alter the expression of key defense-related genes. This has been observed in Arabidopsis infected with *Pseudomonas syringae*, which exhibits both genome-wide hypomethylation and local methylation changes at stress-responsive loci (Pavet et al., 2006). Work with mutants defective in key DNA methylation enzymes, such as MET1, DRM2, and CMT3, further shows altered resistance to pathogens, indicating that proper methylation homeostasis is critical for immune responses (Slaughter et al., 2012, Zhang et al., 2012).

DNA methylation changes induced by stress are not always transient. In some cases, they can be maintained across generations, contributing to transgenerational epigenetic inheritance (Johannes and Schmitz, 2019). For example, under salt stress, hypomethylation of the transposable element *AT5TE35120* near the *CNI1* locus is stably inherited through the male germline and associated with the activation of the long non-coding RNA *CNI1-AS1*. Such cases suggest that stress-responsive methylation changes may play a dual role - enabling immediate gene regulatory flexibility while also contributing to longer-term heritable adaptations (Wibowo et al., 2016, Wibowo et al., 2018).

While many studies have explored DNA methylation changes in plants under stress using Whole-Genome Bisulfite Sequencing (WGBS), their findings are difficult to compare (Table 1). This is primarily due to the use of divergent analysis pipelines, varied thresholds for identifying differentially methylated regions (DMRs), and differing units of analysis, ranging from specific genes to broad genome-wide patterns. These methodological differences limit our ability to identify robust, stress-specific or shared methylation signatures across studies and to draw generalizable conclusions about epigenetic regulation under stress.

**Table 1-.**
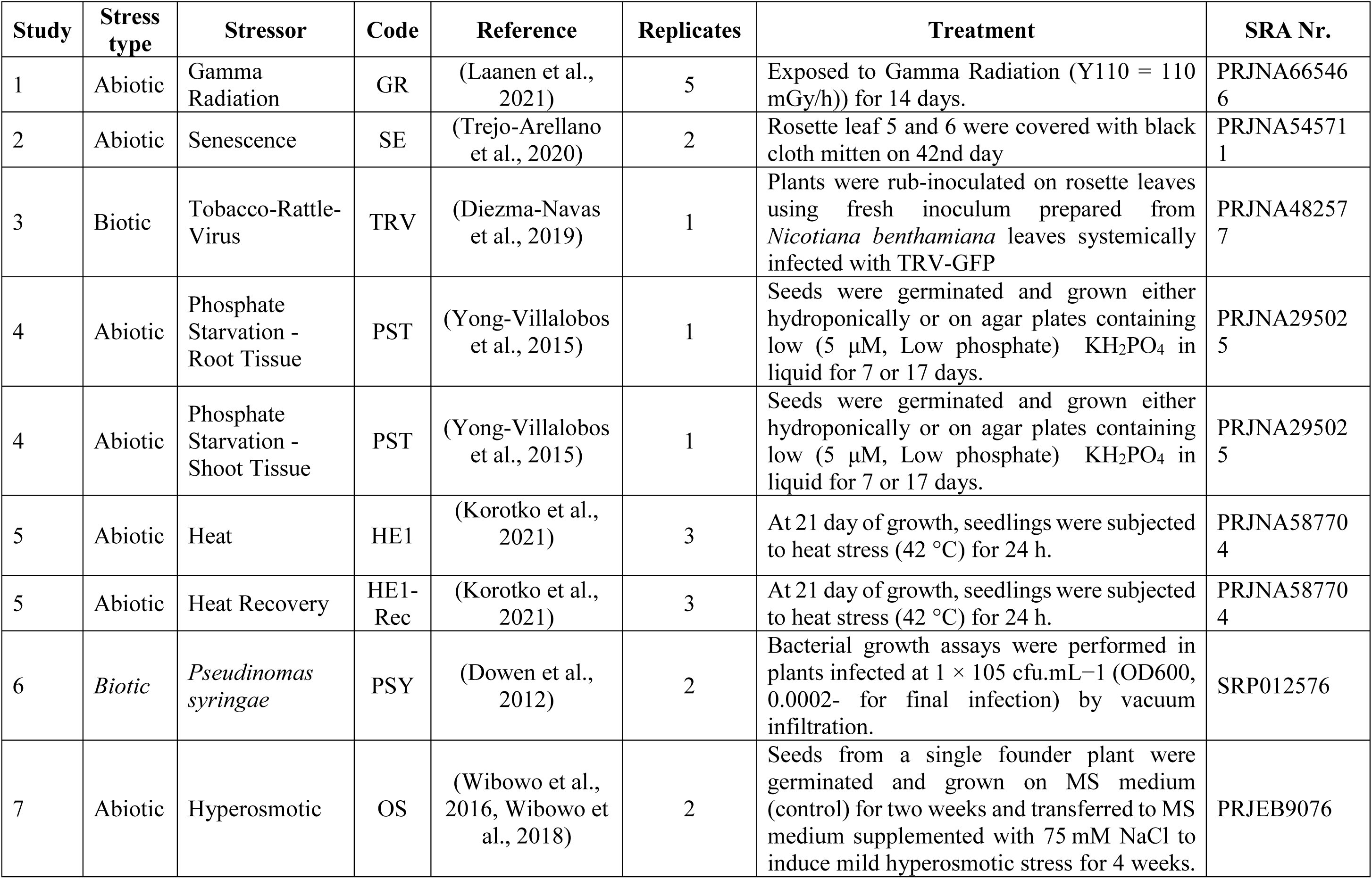
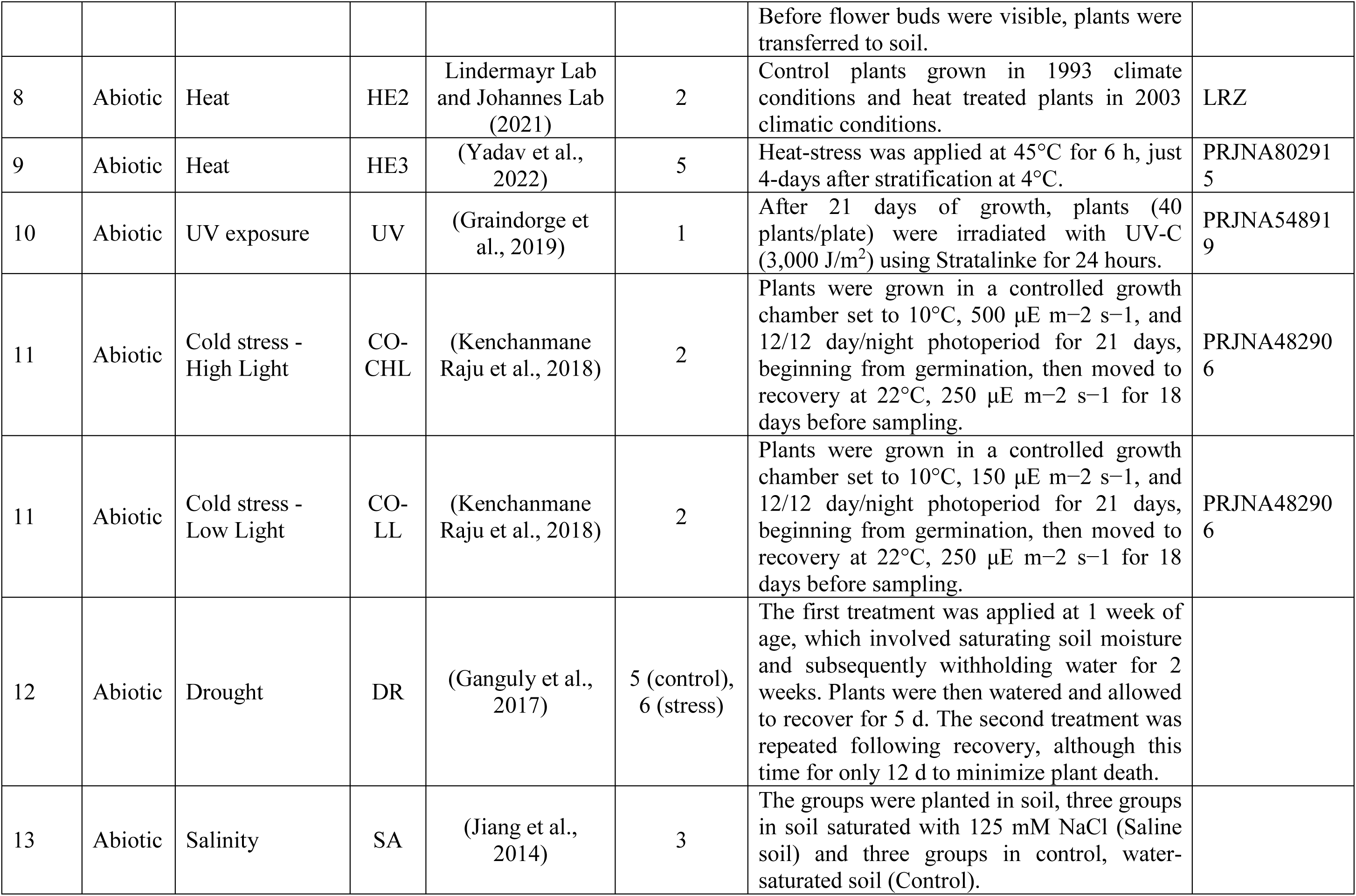
Overview of datasets analyzed for DMR identification under various stress conditions in *Arabidopsis thaliana*.

To overcome this challenge, we conducted an integrative re-analysis of 16 Whole Genome Bisulfite Sequencing (WGBS) datasets from 13 published *Arabidopsis thaliana* studies using a standardized bioinformatic pipeline to identify common and stress-specific methylation responses. By harmonizing DMR detection and annotation across studies, we aim to identify both common and condition-specific methylation responses, with particular focus on stress-regulated genes and TEs. This unified framework provides a more consistent view of the epigenetic landscape associated with stress responses and lays the groundwork for identifying conserved regulatory features and potential targets for improving stress resilience in crops.

## Materials and Methods

### Data Collection: Inclusion and exclusion criteria

WGBS data related to *A. thaliana* was searched and downloaded from NCBI SRA database ((https://www.ncbi.nlm.nih.gov/sra). Following the SRA search, datasets were rigorously screened by reviewing the experimental design of the corresponding studies. Only datasets with a well-defined comparative study between control and stressed (abiotic or biotic) conditions in *A. thaliana* ecotype Col-0 were selected for further analysis. Datasets representing stress responses under both single and multi-generational contexts were included. Conversely, datasets focusing on N^6^-methyladenosine methylation (m^6^A) were excluded. In total, 16 datasets from 13 studies met these criteria and were compiled for further analysis. The associated metadata for each study is presented in Table 1 and Supplementary Figure S1.

### WGBS data processing

WGBS data was processed using the MethylStar pipeline, which integrates tools for read trimming (Trimmomatic), alignment (Bismark) and cytosine-level methylation calling (Methimpute) as described in (Taudt et al., 2018, Shahryary et al., 2020) and https://github.com/jlab-code/MethylStar.

### Methylation level analysis

Global methylation levels were calculated as the average genome methylation across the genome. This quantity reflects the approximate proportion of cytosines that are methylated at the genome-wide scale. Statistical comparisons between control and stress conditions were performed on the global methylation data using an unpaired t-test where more than one replicate was available. For datasets with only a single replicate, the Wilcoxon rank-sum test (Mann-Whitney U test) was applied. Multiple testing adjustments were conducted using the Bonferroni and Holm-Bonferroni methods, with a significance threshold of p< 0.05.

### DMR analysis

DMRs were identified using jDMR (Hazarika et al., 2022) (https://github.com/jlab-code/jDMR). To analyze cytosine clusters, the genome was divided into sliding windows of 500 bp for CG and CHG context and 100 bp for CHH context. Only bins with at least ten cytosines were retained. Methylation region calls were obtained for the sliding windows for all three contexts (CG, CHG, and CHH). Hence, the methylation status of a given region was classified as either unmethylated or methylated. Pairwise comparisons between the control and stress conditions were conducted by generating a DMR matrix containing region calls corresponding to each sample and each bin. Non-epipolymorphic patterns were excluded, and adjacent bins were merged to obtain the final list of DMRs. The number of DMRs within each window were calculated for each study and sequence context, and DMR density was plotted to visualize the distribution of methylation changes across the genome.

### DMR-associated Genes and Transposable Elements (TEs) analysis

Annotation files for TEs, genes and promoters were obtained from TAIR 10, and the intergenic region was delineated using Bedtools. DMR-associated TEs, promoters, intergenic regions and genes were identified for each study considering more than 60% overlap between a DMR and the TE, gene-body, or promoter. Shared DMR-associated genes and TEs among stress conditions were calculated and visualized to highlight similarities or differences across stress conditions. Gene Ontology (GO) enrichment analysis was performed on DMR-associated genes using g:Profiler (FDR = 0.05) ((Raudvere et al., 2019) and https://biit.cs.ut.ee/gprofiler/gost). Thereafter, shared enriched GO terms were compared using the GO Compare graph algorithm for CG, CHG and CHH contexts (Sosa et al., 2023). The distance of each TE to its nearest gene was calculated. These distances were then categorized into four quartiles based on gene-TEs proximity (25% quartile - 1119.25 bp or ∼1.2 kb, 50% quartile 1721 bp or ∼1.7kb, 75% quartile – 2568.75 bp or ∼2.57kb and 100% quartile – 9998 bp or ∼10kb). TEs were further classified into superfamilies, and enrichment analysis of TEs families within each quartile was conducted. The closest 25% quartile of TEs, with the proximity distance of ∼1.2 kb to genes, was selected for further analysis. TE superfamilies within this 25% quartile were extracted, and their enrichment was calculated as the total base pair overlap with DMRs relative to the total TE space. Resulting values were normalized using min-max scaling. Genes located within ∼1.2 kb of these TEs were subjected to GO analysis using g:profiler (FDR = 0.05) to identify potential pathways influenced by TE-associated methylation. All the calculations and visualizations were carried out using R programming and Rstudio, with plot adjustments performed in Inkscape.

### Analysis of spontaneous epimutation rates

To evaluate whether a subset of stress-responsive loci are capable of accumulating transgenerationally stable methylation changes, we analyzed spontaneous CG epimutation rates in wild-type *Arabidopsis thaliana* (Col-0) mutation accumulation (MA) lines using the AlphaBeta R package (v1.10.0) ((Shahryary et al., 2020) and https://github.com/jlab-code/AlphaBeta). We first compiled CG-associated DMRs from multiple stress studies into a unified annotation set. These regions were then assessed in the MA_Col-0 dataset, which includes WGBS data from two independent lineages spanning 17 generations ((Yao, 2023); GEO accession GSE223810). Following the approach of (Hazarika et al., 2022), we estimated the CG methylation gain (α) and loss (β) rates with AlphaBeta. This method builds on earlier modeling frameworks (e.g., (van der Graaf et al., 2015)) and uses multi-generational WGBS data along with known pedigree structures to estimate epimutation rates and their confidence intervals.

## Results

To identify stress-specific and conserved changes in DNA methylation, we performed a comparative analysis of studies (reported in Table 1) to further reveal the function of DNA methylation in plant stress response using a common bioinformatic approach.

### Global DNA methylation levels are stable under stress

Published *A. thaliana* DNA methylation studies using biotic or abiotic stress were extracted from the NCBI-SRA database. In total, 13 publications were identified (Figure 1A) that meet our search criteria (see Methods). These studies included abiotic stressors such as senescence (SE), gamma radiation (GR), phosphate starvation (PST)-root (R) and shoot (S), high osmolarity (OS), UV radiation (UV), cold low light (CO-LL) and high light (CO-CHL), drought (DR), Salinity (SA) and enhanced temperatures (3 studies, HE-1, HE-2 and HE-3 with HE-1Rec, which represents plants that recovered from heat stress) as well as biotic stresses such as Tobacco Rattle Virus (TRV) and *Pseudomonas syringae* (PSY). We re-analyzed the DNA methylation data of these studies using a uniform analysis pipeline (see Methods). A comparison of global CG, CHG and CHH methylation levels between treatment and controls did not reveal statistically significant differences in any of the studies, except in DR (p = 0.003) for CHG context (Figure 1B and Table 2). These results show that global steady-state DNA methylation is robustly maintained in response to biotic or abiotic stress. However, a global analysis potentially masks stress-induced hypo- or hyper-methylated DMRs occurring at specific genomic regions. Their effects may not be detectable in global averages, either because there are too few in numbers, or because hyper- and hypo-methylated DMRs occur with similar frequencies. To assess this we mapped DMRs in each study.

**Figure 1-.**
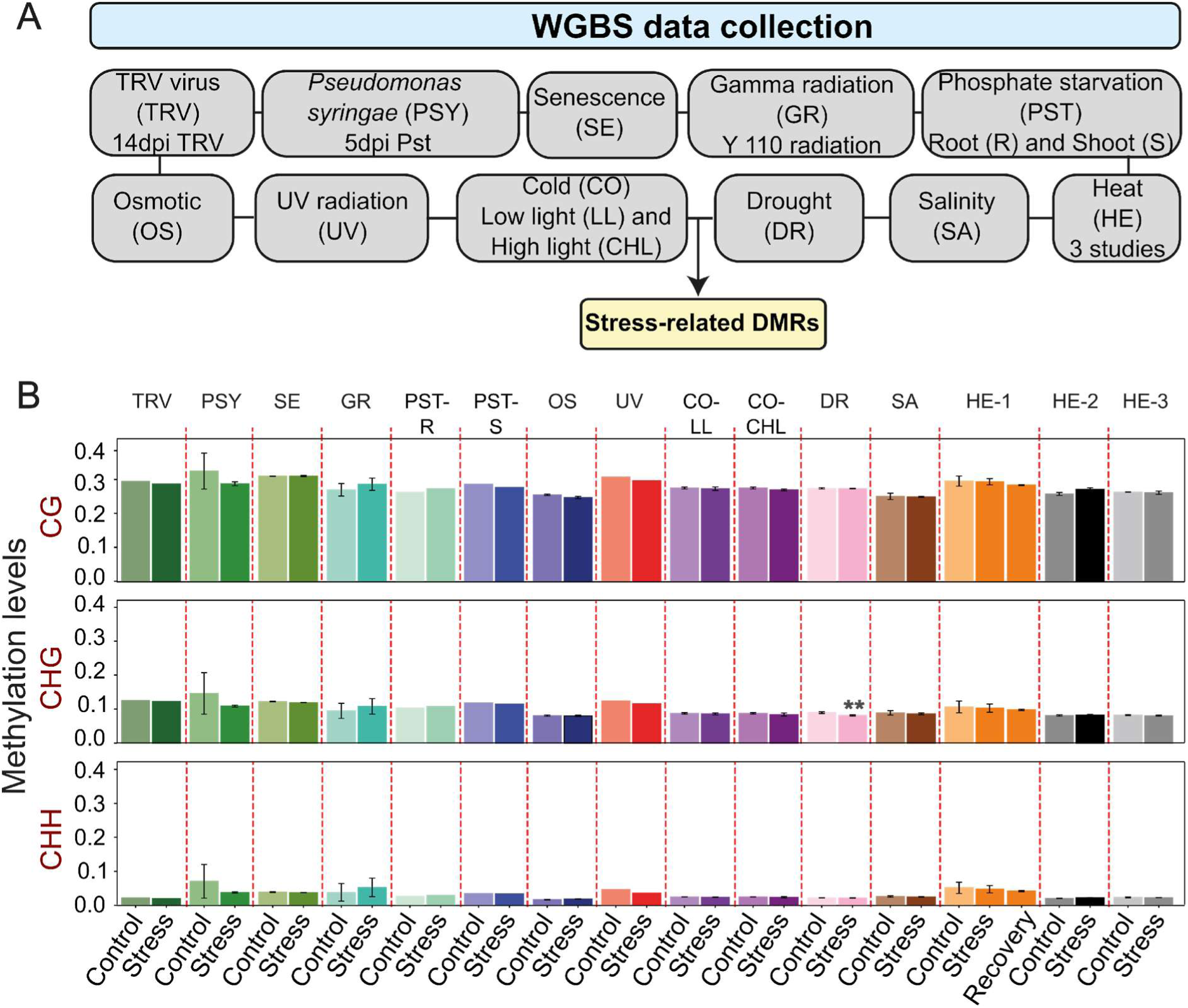
Overview of analyzed studies and methylation levels in different sequence contexts. **(A)** Graphical representation of the experimental conditions analyzed for various studies, including biotic stresses (e.g., *Pseudomonas syringae* infection and TRV virus), abiotic stresses (e.g., UV radiation, drought, salinity, heat, and phosphate starvation), and developmental cues (e.g., senescence). **(B)** Average methylation levels for stress-related methylomes across CG, CHG, and CHH sequence contexts. Each bar represents the mean methylation percentage for the respective context in a specific study, with red dashed lines separating individual experimental conditions. Error bars indicate standard deviations.

**Table 2-.**
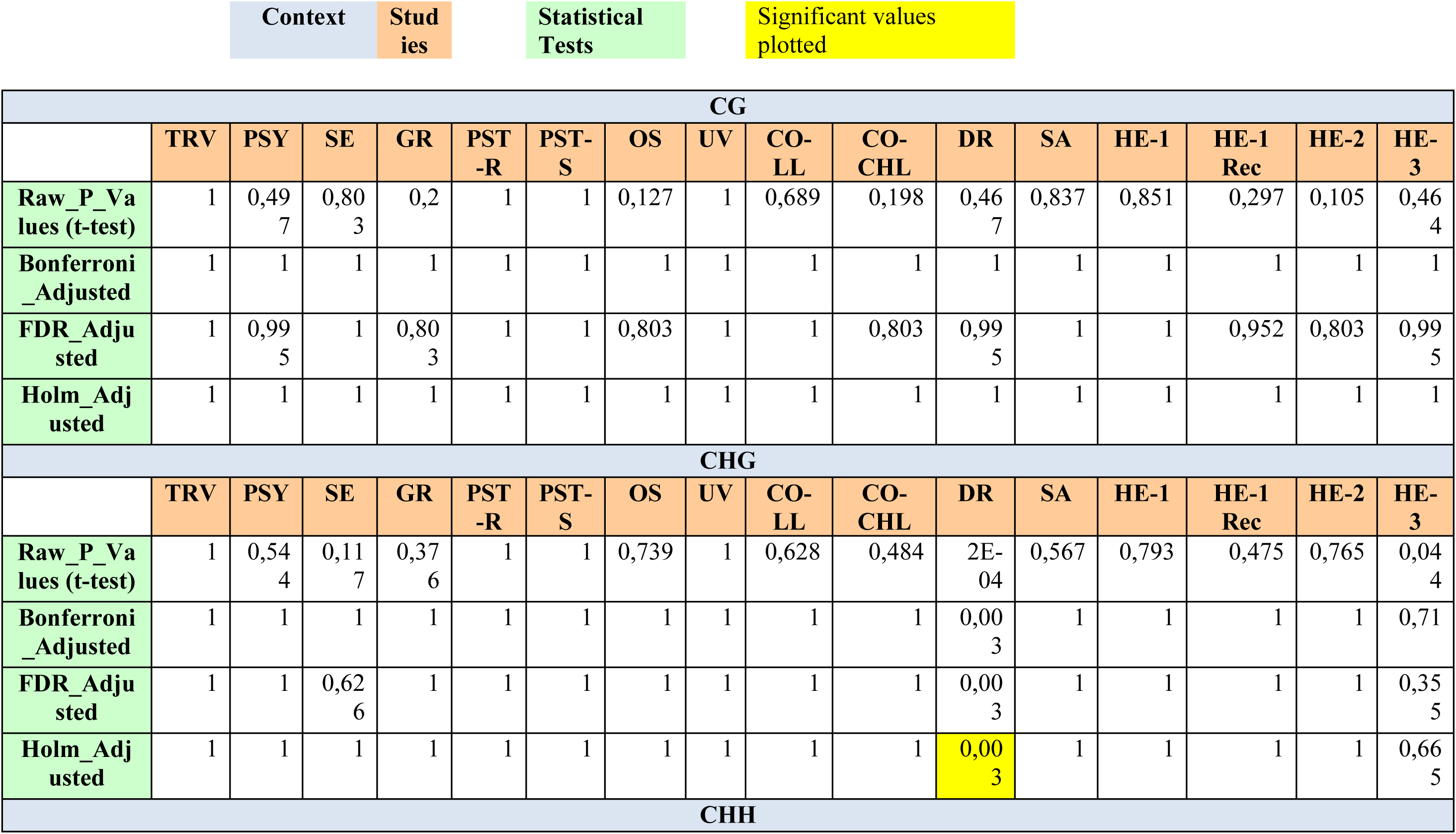
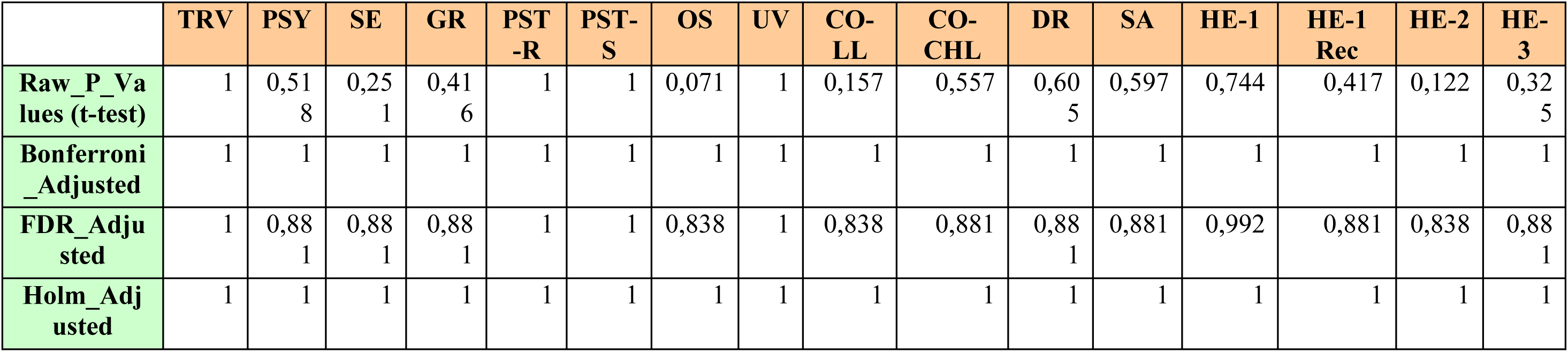
Statistical analysis of global methylation differences across studies for all three contexts – CG, CHG, and CHH. The table presents the results of t-tests comparing control and stress conditions for each study, followed by multiple testing adjustments (Bonferroni, FDR, and Holm methods). Each row represents a statistical test, and columns correspond to studies. Significant values plotted (highlighted in yellow) indicate adjusted p-values below the significance threshold. Contexts (CG, CHG, and CHH) are labeled distinctly to showcase the corresponding methylation analysis

### Differentially Methylated Regions (DMRs) associated with stress conditions

Hyper-methylated DMRs were defined as genomic regions that displayed local methylation increases under stress relative to the controls. Similarly, hypo-methylated DMRs were defined as regions displaying local methylation losses (Figure 2A,B & C and Supplementary Figure S2A,B &C). We identified considerable variation in the total number of DMRs across the different studies, ranging from 52 to ∼13.5k (Figure 2A,B & C). Despite this variability, the proportion of hyper- and hypo-methylated DMRs was very similar for most treatments, with three clear exceptions: PSY showed ∼90% hyper-DMRs in CHG; GR was dominated by hypo-DMRs in CG; and DR by hypo-DMRs in CHG and CHH. In CHH, hyper-DMRs were more frequent under PST, GR, OS, and heat exposures (HE-1, HE-1 Rec, HE-2), whereas CO-LL, CO-CHL, DR, and HE-3 showed more hypo-DMRs. In sum, we did not find a clear trend in the relative frequency of hyper- or hypo-methylation in response to the different stress conditions (Figure 2 and Supplementary Figure S2).

**Figure 2-.**
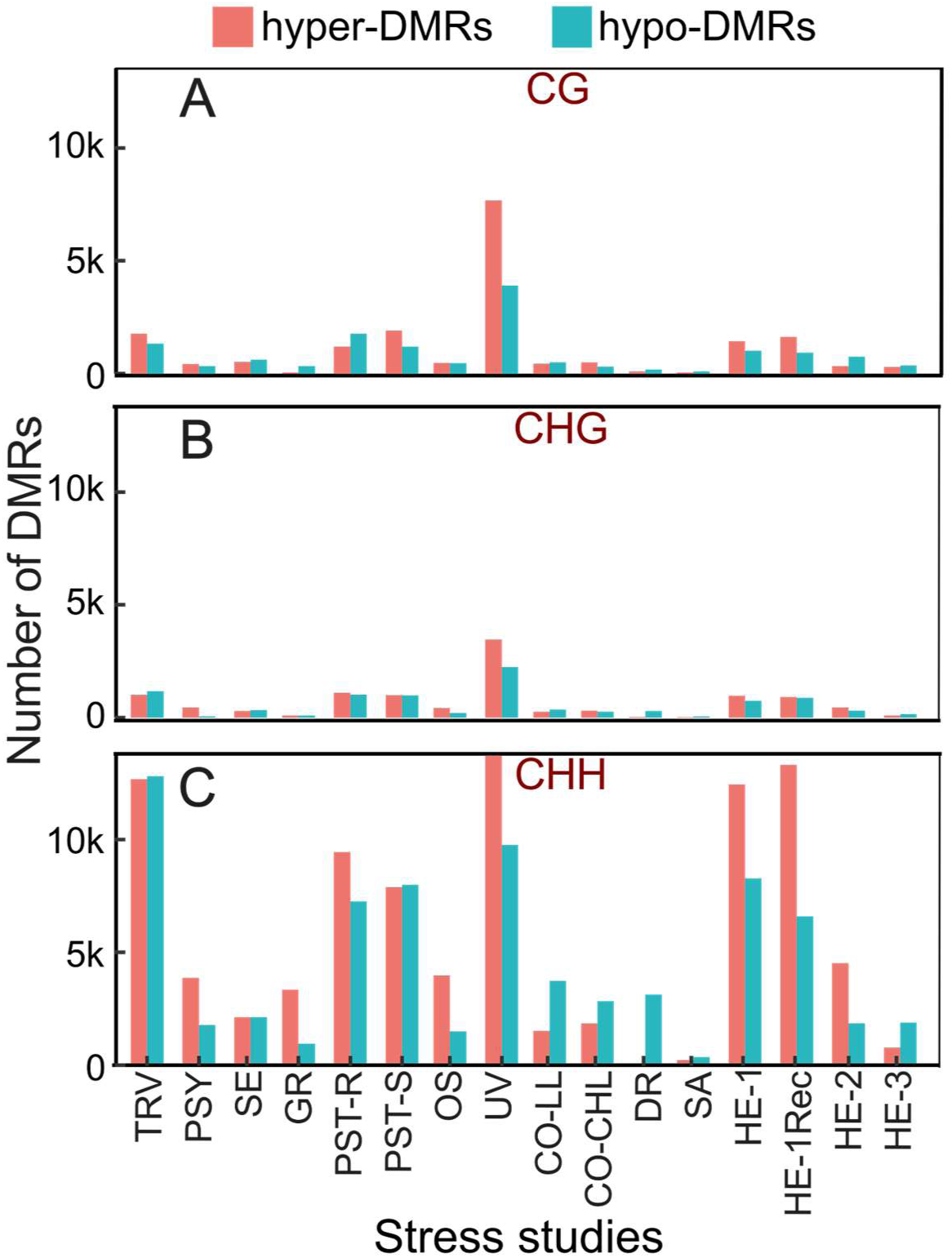
DMRs analysed across different studies. Each bar represents the number of hyper- and hypo-methylated regions for a specific study and sequence context - **(A)** CG **(B)** CHG and **(C)** CHH.

In order to further understand the distribution of DMRs across the *A. thaliana* genome, the DMR density was analyzed in 500kb sliding windows for all five chromosomes (Figure 3). We observed that DMR density across the genome was broadly consistent across studies, with CHG and CHH DMRs clustering mainly in pericentromeric regions of chromosomes, while CG DMRs were more evenly distributed in chromosome arms (Supplementary Figure S3). However, some differences were discernible. For instance, when we compared normalised DMR densities in pericentromeric regions, both CHG and CHH contexts had significantly higher DMR densities than CG contexts on chromosomes 1, 2, 3 and 5 (paired Wilcoxon p<0.05), whereas on chromosome 4 CHG remained above CG (Figure 3 and Supplementary Figure S4 and S5). Finally, splitting DMRs by direction (hyper- vs. hypomethylated) recapitulated the similar genomic distribution patterns (Supplementary Figure S6A & B).

**Figure 3-.**
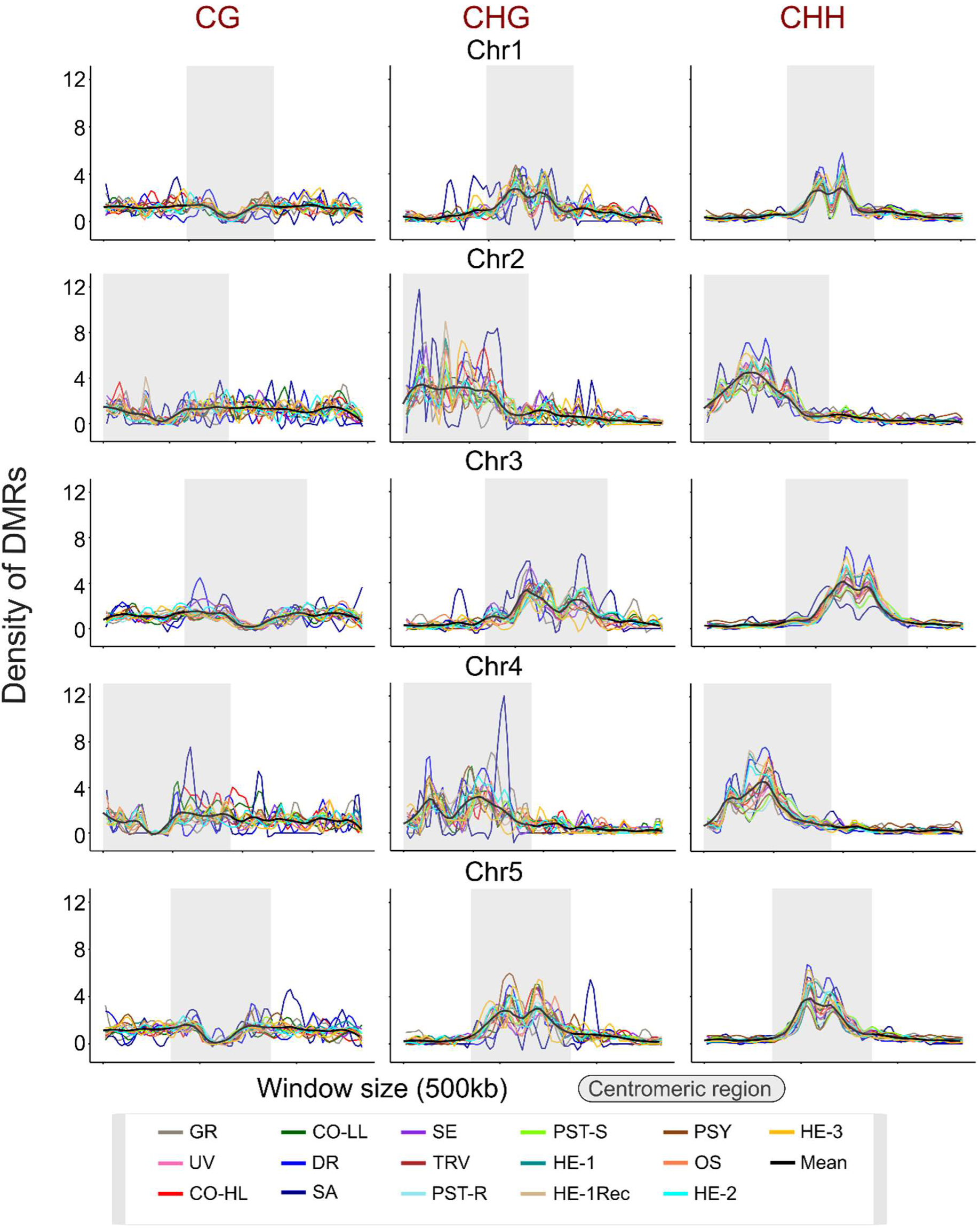
Density of DMRs per genomic window across studies. Densities are shown for three methylation contexts: CG, CHG, and CHH. The gray segment highlights the centromeric region, where DMR density tends to peak in most contexts. Each line represents a different study, as indicated by the legend.

### Annotation-centred DMR analysis

The characteristic density distribution of DMRs suggested that differential methylation in response to stress shows strong preferences for specific genomic features. To examine this in more detail, we annotated all DMRs for their association with genes, TEs, promoters, intergenic regions, and non-coding RNAs (ncRNAs). As expected, most CG-DMRs were associated with genes, whereas CHG-DMRs and CHH-DMRs were predominantly linked to TEs, followed by intergenic regions (Figure 4A, B &C). Only subtle differences in these annotation profiles could be detected across studies. One outlier was PSY, which exhibited an almost 1:1 ratio of CHH-DMRs between TEs and genes. A similar trend was observed when hypermethylated and hypomethylated DMRs were analyzed separately, except in PSY and HE-1-Recovery, where the majority of hypomethylated DMRs were associated with genes in the CHH context rather than with TEs (Supplementary Figure S7A & B). This underscores the nuanced variation in genomic distribution patterns under different stress conditions.

**Figure 4-.**
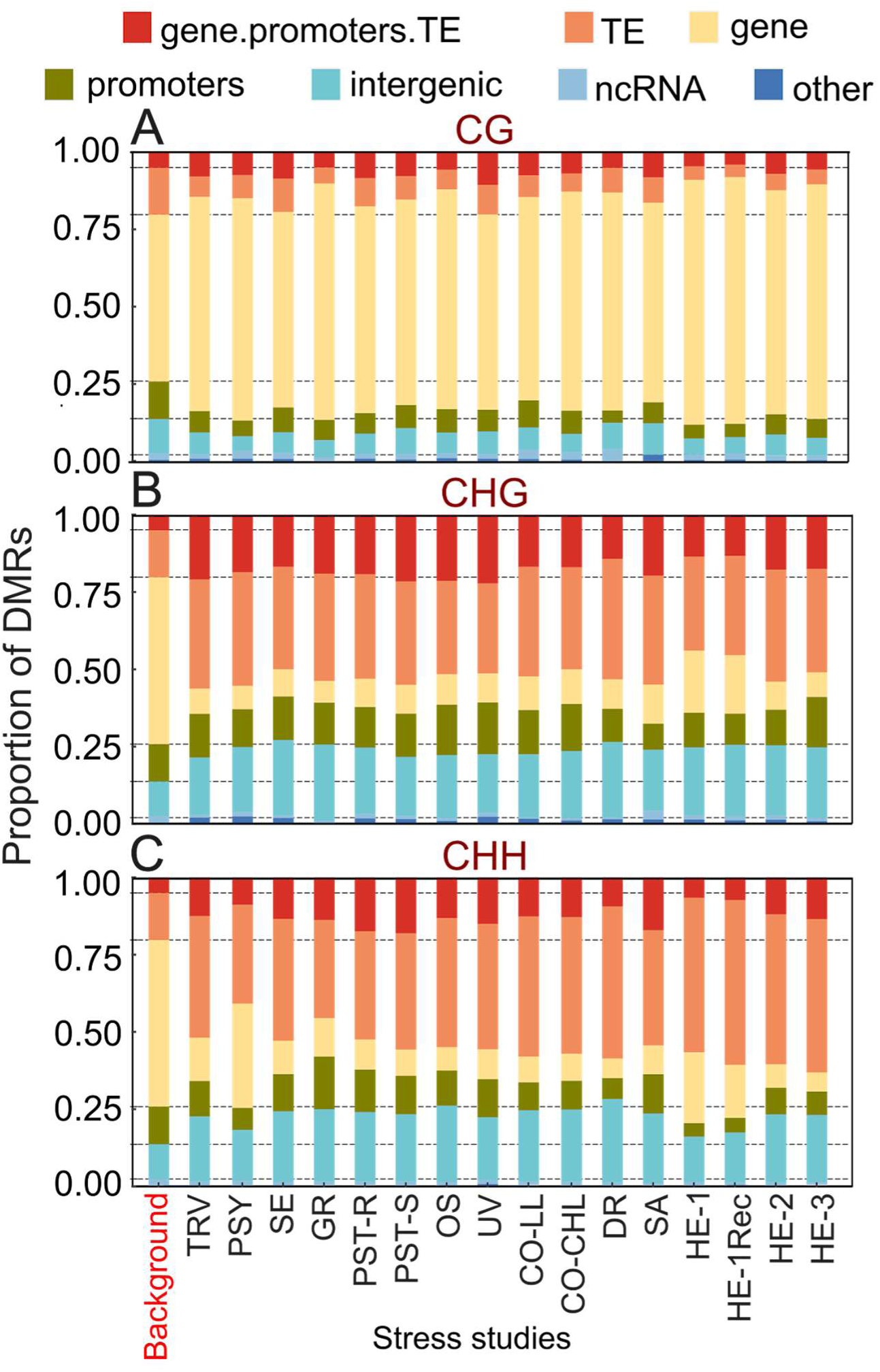
Proportion of DMRs annotated to different genomic features (e.g., gene body, promoter, intergenic regions, etc.) for each study. The three methylation contexts **(A)** CG, **(B)** CHG and **(C)** CHH are analyzed separately. The “Background” bar represents the genomic distribution of features across the entire genome for comparison.

### Gene-based DMR analysis

To identify genes responsive to multiple stress conditions, we analyzed CG-DMRs in genes across studies. In total, we found 10,639 non-redundant genes harboring at least one CG-DMR. Among these, 59.4% (6,320 genes) appeared in at least two studies, and 33.3% (3,542 genes) were shared by three studies (Figure 5A and Supplementary File S1). The number of shared genes decreased with increasing study overlap, with 15 genes present in nine studies and a single gene, *AT1G67500*, found in ten studies. Notably, *AT1G67500* encodes the catalytic subunit of DNA polymerase ζ (REV3), a key player in translesion DNA damage tolerance and UV-damage resistance (TAIR, 2024 and (Sakamoto et al., 2003)).

**Figure 5-.**
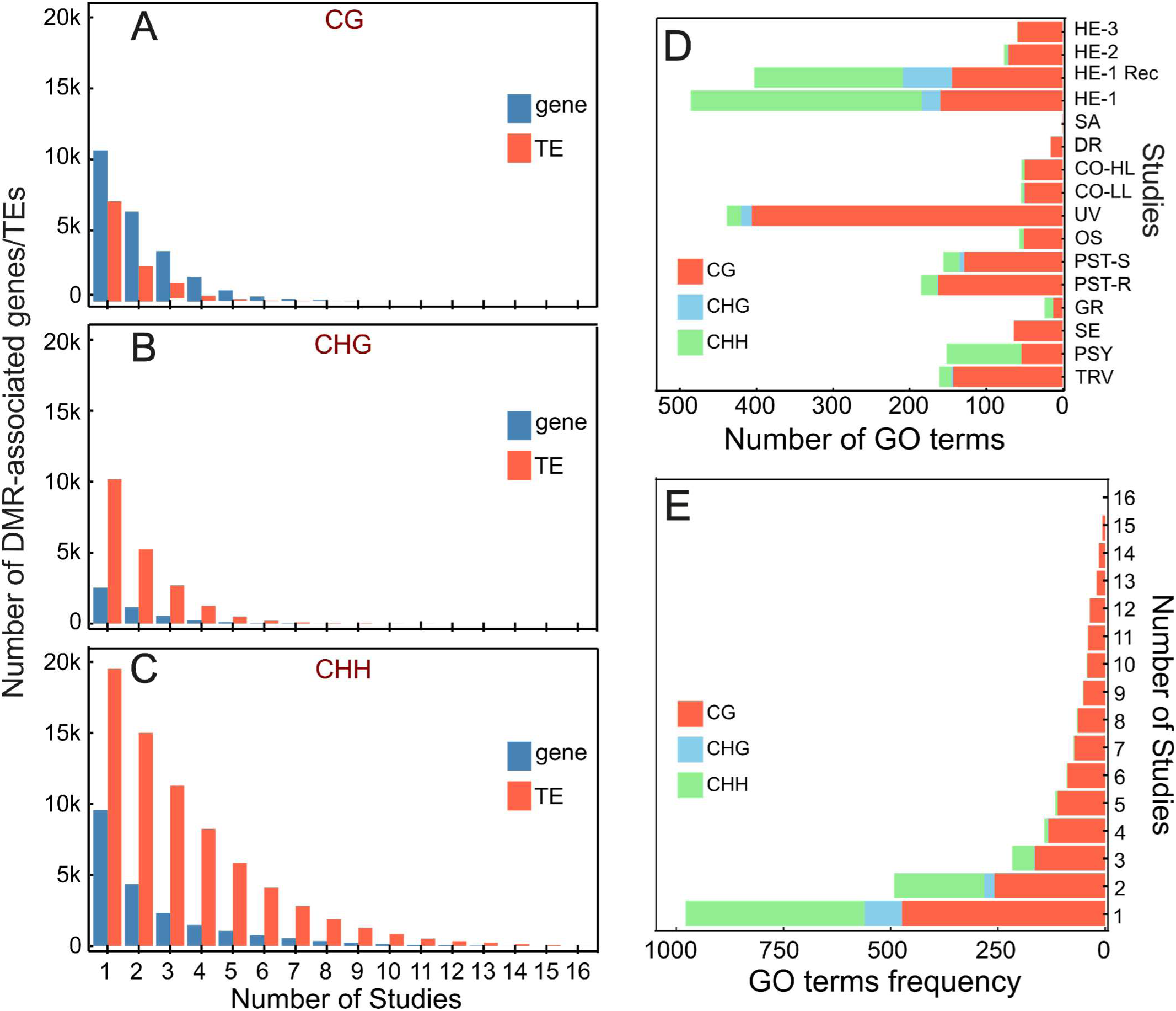
Number of DMR-associated genes and TEs shared across studies. Bar plots show the number of DMR-associated genes (blue) and TEs (red) identified in pairwise comparisons of studies. Each panel corresponds to one of the three methylation contexts: **(A)** CG, **(B)** CHG, and CHH. The x-axis represents the number of studies in which DMR-associated features are shared, while the y-axis indicates the count of shared genes or TEs. **(D)** Number of functional categories (biological processes, BP; molecular function, MF; cellular component, CC) enriched for DMRs in each study. Bars are grouped by study and color-coded to represent the three methylation contexts: CG (orange), CHG (blue), and CHH (green). **(E)** Frequency distribution of enriched Gene Ontology (GO) terms shared across studies for DMR-associated genes in the CG (orange), CHG (blue), and CHH (green) methylation contexts. The y-axis represents the number of GO terms, and the x-axis indicates the number of studies sharing the enriched terms across the three GO categories: BP, MF and CC.

In comparison, CHG-DMR yielded 2,540 unique annotated genes. Among these, 45.7% (1,159 genes) were shared across at least two studies, and 21.6% (548 genes) were common among three studies (Figure 5B and Supplementary File S1). Genes such as *AT3G45140* (chloroplast lipoxygenase LOX2) and *AT1G33950* (Immune-associated nucleotide (IAN)-binding protein 7) appeared in ten and eight studies, respectively (TAIR, 2024). LOX2 has documented roles in jasmonate signaling and abiotic stress - particularly wounding, UV, and osmotic stress - mediated via singlet oxygen production in roots (Chauvin et al., 2013, Chen et al., 2021, Singh et al., 2022). IAN family genes are typically upregulated during plant– pathogen interactions and are linked to immune signaling. While IAN7 has not been extensively characterized, it is classified in the AIG1 family - known for responses to pathogen infection - and shows differential expression during immune-related processes (Lv et al., 2019). In the CHH context, 9,589 annotated genes overlapped with DMRs. Among these, 45.4% (4,350 genes) were shared among at least two studies, and 24.3% (2,326 genes) were common among three studies (Figure 5C; Supplementary File S1). Interestingly, four genes were shared across 15 studies: *AT2G02635* (unknown function), *AT2G07240* (cysteine-type peptidase), *AT2G10625* (Ulp1 protease family), and *AT2G11851* (unknown function), and one gene (*AT2G07240*-cysteine-type peptidase) was shared across all 16 studies, suggesting a possible role in stress resilience via proteolytic processing (Brocklehurst et al., 2018).

To assess whether DMR-associated genes identified in each sequence context are part of a conserved regulatory response, we performed a bootstrap analysis using all annotated Arabidopsis genes as a background ((Berardini et al., 2015) and Araport11). We found that the overlapping set of DMR-associated genes was non-random across studies (Bootstrap p<0.0001), indicating that DNA methylation responses to diverse stressors converge on a set of shared methylation-responsive genes (Supplementary Figure S9). A similar pattern was observed when analyzing hyper- and hypomethylated DMRs separately (Supplementary Figure S8, S10 and Supplementary File S9 & S10).

### Functional enrichment of DMR-associated genes

To investigate biological processes potentially regulated by differential methylation, we used gProfiler to perform Gene Ontology (GO) term enrichment analysis on the differentially methylated genes from each stress study. Various enriched GO terms (functional categories) were identified for all the studies in the CG context. However, for some studies in the CHG (GR, SE, PSY, OS, HE-3, CO-HL, DR and SA) and CHH (SE, DR and SA) contexts, no significantly enriched GO terms were detected. In general, more enriched GO terms were found in the CG context due to the greater number of genes identified in this context (Figure 5D). Specifically, the highest number of enriched GO terms were observed in the HE-1 and HE-1 Recovery studies.

To analyze overlapping stress responses, common GO terms among studies were identified. In the CG context, 474 enriched GO terms were found across all studies, with 258 shared between at least two studies and 164 among three studies. Fifteen studies shared seven common GO terms, primarily related to binding processes such as anion or ATP binding (Figure 5D and Supplementary File S2). In the CHG context, 87 enriched GO terms were identified, but only a few were shared across multiple studies, with cell wall structure and extracellular region being the most commonly enriched terms (Figure 5D and Supplementary File S3). In the CHH context, 417 enriched GO terms were identified, with common terms associated with cell wall structure and hydrolase enzyme activity (Figure 5D and Supplementary File S4).

Further stratification of genes into gbM versus non-gbM genes revealed distinctive enrichment patterns. In general, non-gbM genes exhibited more enriched GO terms across CHG, and CHH contexts compared to CG context, suggesting these genes play a more active role in stress-responsive regulatory networks (Supplementary Figure S12A, B and C). This aligns with previous findings indicating that non-gbM genes, which are generally lowly methylated, are more responsive to epigenetic alterations under stress conditions (Horvath et al., 2019, Muyle et al., 2022, Zhang et al., 2024).

A Fisher exact test further supported this differential pattern of enrichment (one-sided Fisher’s exact test; p<0.05). Non-gbM genes in the CG context were particularly enriched for GO terms (top 12) related to cellular organization and metabolic processes. In contrast, enriched GO terms among gbM genes were primarily associated with basic cellular processes such as enzymatic activities (e.g., kinase activity or phosphorylation), consistent with their putative housekeeping roles. For CHH context, significant non-gbM GO terms were predominantly associated with cell wall structure, extracellular regions, and hydrolase enzyme activity. These results suggest that differential methylation events, especially in non-gbM genes, target specific stress-responsive pathways (Supplementary Figure S12D, E and F). Together, these findings imply that, despite variations across stress types and methylation contexts, conserved biological mechanisms involving non-gbM genes may significantly contribute to stress adaptation.

We used a network-based analysis to further visualize the relationships between stress conditions and enriched GO terms (Figure 6). In the CG context, UV stress showed the highest number of enriched GO terms, especially those related to binding (e.g., ATP, adenyl nucleotide, anion, carbohydrate) and enzymatic activities (ATP hydrolysis, catalytic activity, transferase activity). TRV, PST-R, PST-S, and HE-1 also shared many of these terms, whereas SA and DR exhibited limited GO terms, with SA enriching only ATP-dependent activity and DR highlighting GABA biosynthesis and cell-surface receptor signaling (Figure 6A and Supplementary File S5).

**Figure 6-.**
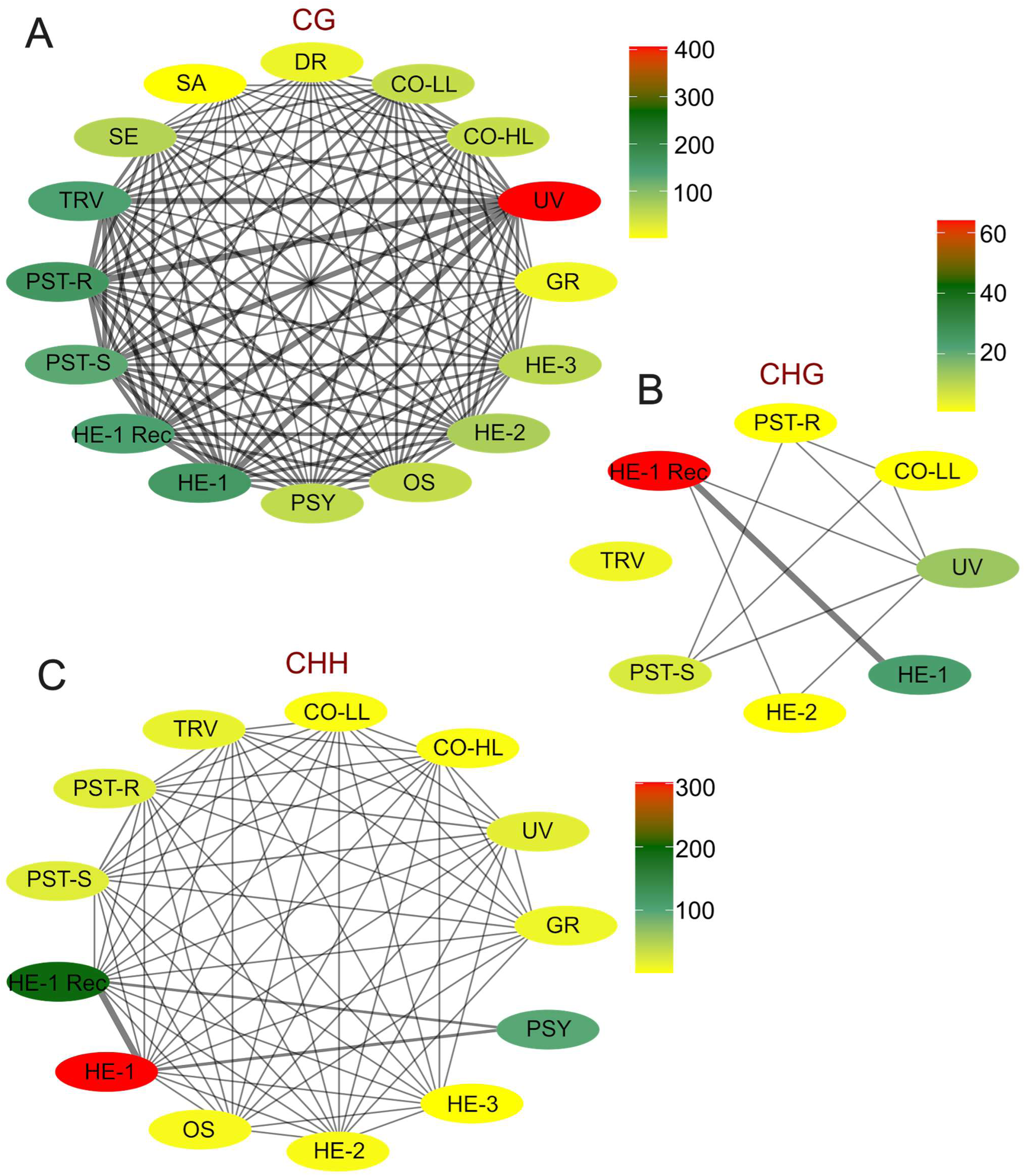
Network graphs illustrating the relationships between studies based on enriched GO categories. The node colors correspond to the number of GO categories enriched in each study (green: fewer categories; red: more categories). The edge widths indicate the number of shared GO terms between studies, with thicker edges representing greater overlap. **(A)** CG context **(B)** CHG context **(C)** CHH context.

In the CHG context, HE-1 and HE-1 Recovery had the most enriched terms - anchoring and cell-wall junctions and structural molecule activities - reflecting cell-wall modifications during heat stress. These findings align with earlier research underscoring structural integrity and cell wall modifications as critical during heat stress adaptation (Le Gall et al., 2015). CO-LL, PST-R, PST-S, and UV shared structural cell-wall constituents, while TRV showed no overlap (Figure 6B and Supplementary File S5). In CHH, HE-1, HE-1 Recovery, and PSY were prominent for cell-wall structure, cellular organization, and enzymatic activities; PSY, bacterial and external stimulus responses and UV, DNA polymerase activity (Figure 6C and Supplementary File S5). Overall, these GO-term enrichments reveal both shared and stress-specific methylation-driven processes - particularly those governing binding functions, cell-wall integrity, and enzymatic pathways - highlighting conserved molecular mechanisms that underpin plant adaptation to diverse stress conditions.

### TE-based DMR analysis

Most plant genomes contain a significant proportion of TEs, which play a crucial role in genome evolution (Lisch et al., 2013, Secco et al., 2015). Environmental changes can activate specific TE families, leading to new insertions and altered gene regulatory landscapes (Dubin et al., 2018). TE activity is tightly controlled by DNA methylation, which typically maintains them in a silenced state (Chuong et al., 2017). However, TEs are not merely genomic parasites - they can also function as gene expression modulators in response to environmental cues. Many TEs contain cis-regulatory elements, such as stress-responsive motifs, that allow them to act as environmental sensors. For example, the heat-responsive TE ONSEN becomes transcriptionally activated under elevated temperatures and can influence the expression of nearby genes (Lisch et al., 2013, Pietzenuk et al., 2016, Chuong et al., 2017). Such environmentally sensitive TEs may be co-opted by the host genome to fine-tune gene expression under stress.

To identify differentially methylated TEs shared across stress studies, we collated 7,051 CG-DMR–associated TEs and found that 2,489 (35.3 %) were shared by at least two studies, dropping to 1,045 (14.8 %) in three studies (Figure 5A and Supplementary File S1). Notably, four TEs (*AT1TE48745/LTR/Gypsy*, *AT1TE50315/LTR/Gypsy*, *AT1TE50375/LTR/Gypsy*, *AT1TE50810/LTR/Gypsy*) recurred in seven studies, and *AT1TE50315* appeared in eight - consistent with previous reports that a core set of TE insertions is stably maintained and often differentially methylated across diverse Arabidopsis accessions and conditions (Stuart et al., 2016, Arora et al., 2022).

In the CHG context, 10,179 CHG-DMR–associated TEs were detected, of which 5,229 (51.4 %) were shared by ≥2 studies and 2,702 (26.5 %) by ≥3 (Figure 5B). Three CHG-TEs (*AT2TE22065/LTR/Gypsy, AT3TE51685/*LTR/Gypsy*, AT5TE35915/*RC/Helitron) were present in ten studies, and two (*AT2TE22065, AT3TE51685*) in eleven (Figure 5B and Supplementary File S1). The CHH context showed the highest level of sharing: of 19,504 CHH-DMR–associated TEs, 14,997 (76.9 %) occurred in ≥2 studies and 11,300 (57.9 %) in ≥3 (Figure 5C). Eight CHH-TEs (e.g., *AT2TE00015/*LTR/Gypsy*, AT2TE10385/*RC/Helitron*, AT2TE11515/*DNA/MuDR*, AT2TE16220/*RC/Helitron*, AT2TE17095/*RC/Helitron*, AT4TE17970/*RC/Helitron*, AT5TE45920/*RC/Helitron*, AT5TE47895/*RC/Helitron) were common to all 16 studies (Figure 5C and Supplementary File S1).

To assess whether DMR-associated TEs identified in each sequence context are part of a conserved regulatory response, we performed a bootstrap analysis using the full complement of ∼32,000 annotated Arabidopsis TEs (Supplementary Figure S9) (Quesneville, 2020). We found that the overlapping set of DMR-associated TEs was non-random across studies (Bootstrap pvalue < 0.0001), indicating that DNA methylation responses to diverse stressors converge on a set of shared methylation-responsive TEs (Supplementary Figure S9). A similar pattern was observed when analyzing hyper- and hypomethylated DMRs separately (Supplementary Figure S8, S11 and Supplementary File S9 & S10).

**Figure 7-.**
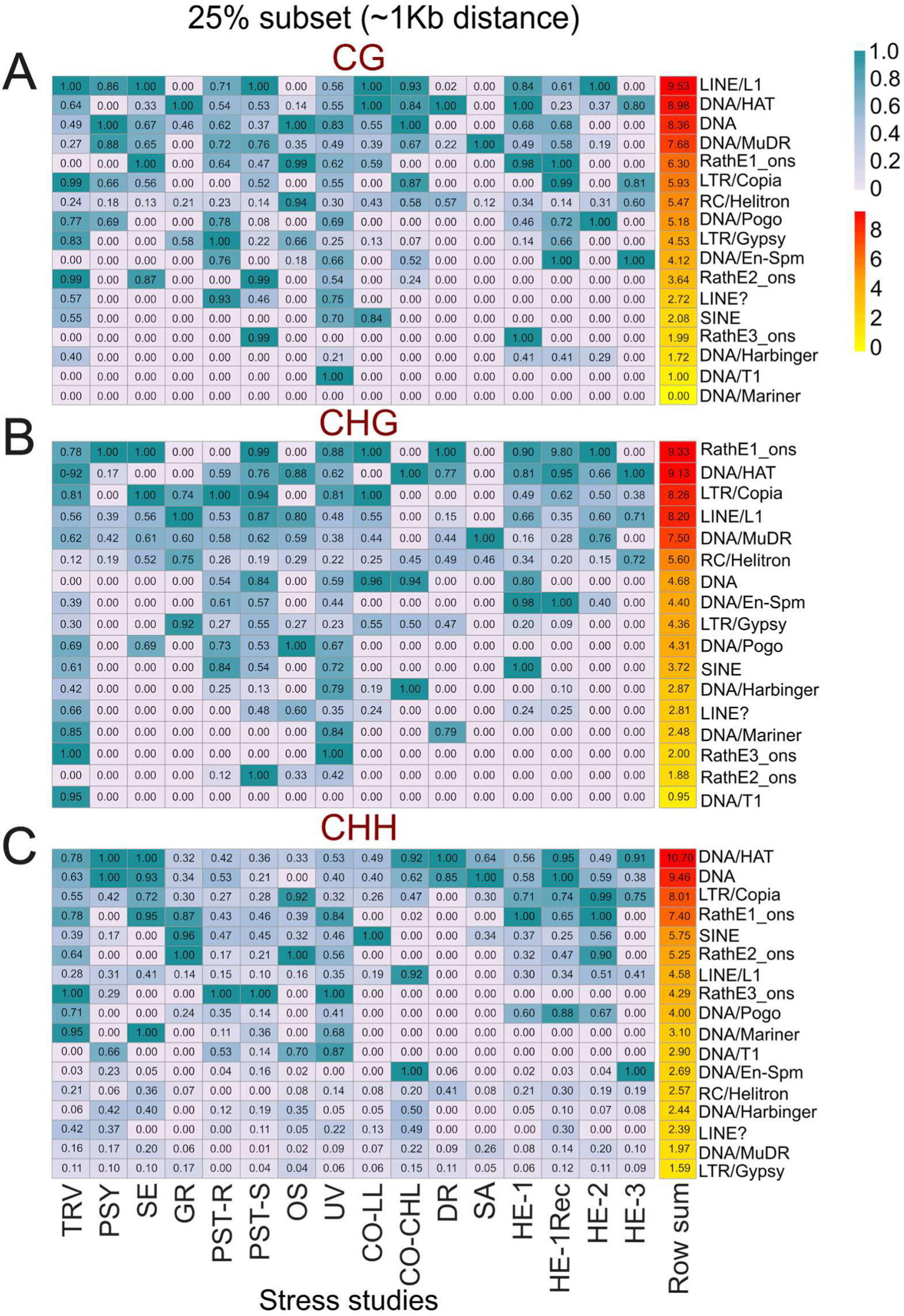
Heatmap showing TE superfamilies located within 25% (approximately 1.2 kb) of DMR-associated genes, sorted in descending order of their enrichment for **(A)** CG, **(B)** CHG and **(C)** CHH contexts. The color gradient reflects the level of enrichment, with red indicating higher enrichment and blue indicating lower enrichment.

### Gene-TE proximal DMR analysis (+ GO analysis of genes in close proximity to TEs)

To determine the relationship between genes and TEs in close proximity, we analyzed the density of TEs relative to their distance from nearby genes. The results (Supplementary Figure S13A) indicated that 25% of all TEs were located within 1.2 kb of the nearest gene, both upstream and downstream, while 50% were within 1.7 kb. We further examined TE superfamilies for different distance subsets (25%, 50%, 75%, and 100%) and calculated the proportion of sequence space occupied by each superfamily (Supplementary Figure S13B). Among TEs located near genes, RC/Helitron was the predominant superfamily, accounting for 46% of the total TEs in the 25% subset and increasing to 61% in the 75% subset. However, when considering all TEs (100% subset), the proportion of RC/Helitron decreased to 32%. Overall, the majority of differentially methylated TEs near genes belonged to the RC/Helitron, DNA/MuDR, LTR/Copia, and LTR/Gypsy superfamilies.

To investigate the enrichment of differentially methylated TEs near genes, we restricted our analysis to all TEs located within 1.2 kb (25% subset) of an annotated gene. For each TE superfamily and each stress study, we then calculated the proportion of these proximal TEs overlapping DMRs in CG, CHG, and CHH sequence contexts. To assess statistical significance, we performed 10,000 permutations of the DMR-overlap labels across that same fixed set of proximal TEs, yielding a null distribution of enrichment for each family. In the CG context, the highest enrichment (Δ% true overlap vs. the permuted overlap) was seen for LTR/Copia in TRV, PST-R, UV and CO-CHL; RC/Helitron in PSY, PST-R and UV; DNA/MuDR in PST-R and UV; DNA/Pogo in HE-1 recovery; and DNA/HAT in HE-3 (p < 0.01 for each). This indicates that, among TEs within 1.2 kb of genes, these superfamilies are significantly over-represented among those overlapping CG-DMRs. To our knowledge, no prior studies have linked RC/Helitron, DNA/Pogo, DNA/MuDR, or DNA/HAT elements to stress responses in Arabidopsis. By contrast, (Ito et al., 2011) previously reported heat-activated mobilization of an ONSEN LTR/Copia element, which we did not observe in our CG-DMR data (Supplementary Figure S14 and S15).

In the CHG context, DNA/MuDR, LTR/Copia, and RC/Helitron superfamilies were each significantly over-represented among gene-proximal CHG-DMRs (all p < 0.05). In the CHH context, we observed a distinct pattern: DNA/HAT, DNA/En-Spm, DNA/MuDR, LTR/Gypsy, and RC/Helitron were all significantly enriched for CHH-DMRs (p < 0.05).

Strikingly, across all three sequence contexts the families DNA/HAT, DNA/MuDR, and RC/Helitron comprised a conserved core of stress-responsive elements. By contrast, other superfamilies such as DNA/Harbinger, DNA/T1, and LINE/L1 showed significant enrichment only under specific treatments (e.g. DNA/Harbinger in UV for CHH; LINE/L1 in HE-2 for CHH), underscoring both a universal TE–methylation signature and treatment-specific dynamics.

We next asked whether genes bearing nearby TE-DMRs might have documented stress-related functions. Across all three sequence contexts, GO-enrichment of proximal genes highlighted core processes-binding and enzymatic activities; transport; reproduction/development; defense; cell-wall biogenesis; and, notably, chromatin remodeling in CG and ncRNA regulation in CHH (Figure 8A–C; Supplementary Files S6–S8).

**Figure 8-.**
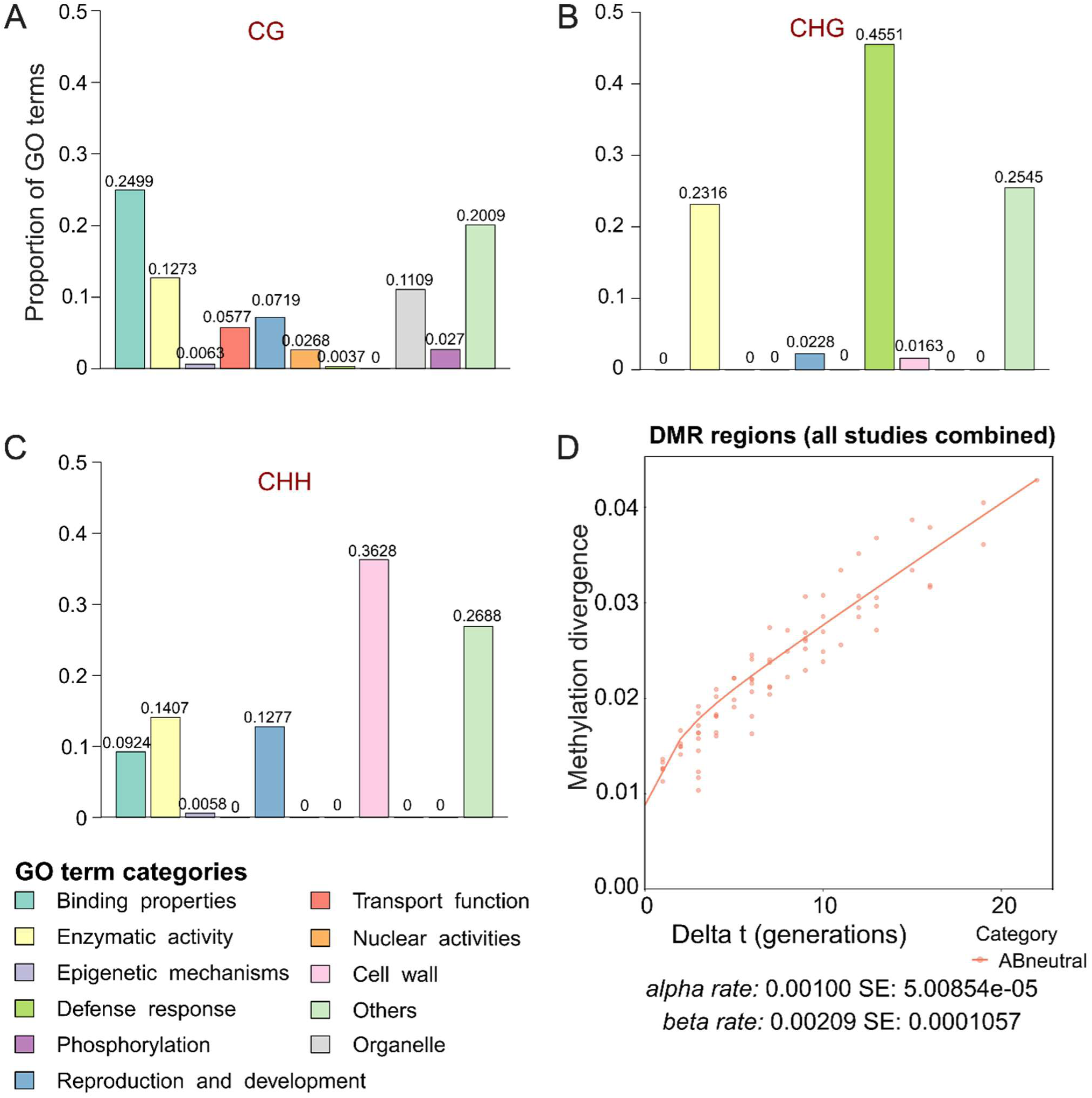
Histograms showing the results of functional enrichment analysis for genes located in 1 kbp close proximity to differentially methylated TEs. The x-axis represents functional categories or GO terms, and the y-axis represents the number of genes associated with each category. **(A)** Genes near TEs in the CG methylation context. **(B)** Genes near TEs in the CHG methylation context. **(C)** Genes near TEs in the CHH methylation context. **(D)** AlphaBeta analysis illustrates the relationship between 5-methylcytosine (5mC) divergence and delta time, describing the accumulation of spontaneous epimutations over successive generations. The x-axis represents delta time, while the y-axis indicates methylation divergence. Data points reflect observed values, and the trendline highlights the spontaneous epimutation accumulation pattern.

In the CG context, one of the top candidates was *AT3G44570*, which encodes a retrotransposon ORF1 protein that assembles with L1 RNA into a ribonucleoprotein particle essential for L1 transposition (Martin et al., 2006). In the CHG context, *WRKY42* emerged in five independent studies; in rice its ortholog confers resistance to the fungal pathogen *Magnaporthe oryzae* (Phukan et al., 2016). Finally, in the CHH context the most recurrent hit was *AT4G16860* (*RPP6*) found in 13 of 16 stress datasets, including TRV and PSY - consistent with its established role in Arabidopsis resistance to downy mildew (van der Biezen et al., 2002) (Supplementary File S12). Together, these examples illustrate how TE-proximal methylation changes can pinpoint genes with previously validated roles in stress and immune responses.

### Potential for transgenerational inheritance of stress-induced CG methylation

While stress-induced DNA methylation changes are often considered transient - typically reprogrammed during reproduction - emerging evidence suggests that some of these changes, particularly in the CG context, may persist across generations (Johannes and Schmitz, 2019). This raises the possibility that stress could contribute to long-term, heritable epigenetic variation. As our DMR analysis showed, most stress-responsive methylation changes occur in TEs, which are targeted by the RNA-directed DNA methylation (RdDM) pathway (Law et al., 2010, Law et al., 2013, Matzke and Mosher, 2014). This pathway reinforces methylation during gametogenesis, thereby resetting most environmentally induced changes and preventing their transgenerational inheritance. However, not all genomic regions are under strong RdDM control. Recent studies have shown that CG sites outside of RdDM targets can undergo spontaneous epimutation; that is, heritable stochastic gains or losses of methylation that arise during imperfect methylation maintenance in mitosis or meiosis (Becker et al., 2011, Schmitz et al., 2011, Schmitz and Ecker, 2012, van der Graaf et al., 2015, Hofmeister et al., 2020, Denkena et al., 2021, Hazarika et al., 2022, Yao et al., 2023). These CG epimutations, unlike their CHG and CHH counterparts, can accumulate over generations at rates far exceeding those of genetic mutations, offering a substrate for long-term epigenetic divergence. We hypothesized that CG DMRs identified in stress-response studies may overlap with genomic regions prone to spontaneous epimutations, providing a mechanistic link between stress-induced methylation and stable, heritable changes in gene regulation. Although direct transgenerational data are lacking for the stress datasets we analyzed, we assessed the potential stability of stress-responsive CG DMRs by examining their behavior in Arabidopsis wild-type mutation accumulation (MA) lines (Yao et al., 2023). These lines, derived from single founders and propagated by selfing for 17 generations, provide a robust system for studying epimutation accumulation and estimating epimutation rates. We focused our analysis on CG DMRs identified in the comparative stress dataset and observed that CG sites within these regions accumulate epimutations over successive generations (Figure 8D). This finding indicates that methylation changes - whether arising spontaneously or triggered by environmental stress - can persist over time, serving as a potential interface between external stress cues and long-term epigenetic inheritance. Such stability suggests a mechanism by which environmental conditions may leave a heritable molecular imprint, potentially influencing adaptive trajectories across generations.

## Discussion

Our reanalysis of *Arabidopsis thaliana* methylomes under diverse biotic and abiotic stresses highlighted critical insights into plant epigenetic dynamics and their implications for adaptive responses. The observed global stability in steady-state methylation levels aligned with previous findings indicating that genome-wide methylation patterns are typically robust, suggesting effective homeostatic mechanisms in plants (Kawakatsu et al., 2016, Bewick and Schmitz, 2017). However, localized changes at DMRs revealed highly context-dependent methylation dynamics, emphasizing the nuanced role of DNA methylation in stress adaptation.

The divergence observed in DMR patterns across different stress contexts, notably the hypermethylation of CHG DMRs under biotic stress (e.g., *Pseudomonas syringae*) and hypomethylation under drought and gamma irradiation stress, paralleled previous studies demonstrating the stress-specific activation of methylation pathways (Dowen et al., 2012, Ganguly et al., 2017).

Chromosomal localization patterns of DMRs, particularly the enrichment of CHG-context DMRs within pericentromeric heterochromatin, confirmed earlier studies that linked heterochromatic methylation to transcriptional silencing and genome stability under stress (Zemach et al., 2013, Stroud et al., 2014).

The substantial overlap of stress-responsive DMR-associated TEs across treatments, particularly involving TE superfamilies such as RC/Helitron, DNA/MuDR, and DNA/HAT, aligned with previous evidence linking TEs to rapid genomic adaptation via methylation-mediated transcriptional regulation (Lisch et al., 2013, Quadrana et al., 2016, Stuart et al., 2016). Some studies have documented the impact of differentially methylated TEs on gene expression (Dubin et al., 2018, Deneweth et al., 2022). For instance, the OS study identified methylation changes in the TE *AT5TE35120*, which is linked to *CNI1* expression across generations exposed to salt stress (Wibowo et al., 2016, Wibowo et al., 2018). Our analysis corroborated these findings, revealing hypomethylation of *AT5TE35120* in the CG context, supporting the hypothesis that CNI1-AS1 long non-coding RNA (lncRNA) expression is associated with the demethylation of this region. These TE-mediated regulatory networks potentially facilitated rapid epigenetic adjustments to environmental challenges, influencing gene expression patterns necessary for plant adaptation.

While the prevailing view holds that stress-induced epigenetic changes are transient (Lämke and Bäurle, 2017, Zhang et al., 2018), our findings suggest that a small subset of CG methylation changes acquired during stress have the potential for long-term inheritance. Although we did not observe this inheritance directly, we found that stress-induced CG DMRs overlap with cytosines known to accumulate stable, stochastic, and heritable epimutations. Prior work with MA lines has shown that methylation gains and losses in these regions are remarkably stable over many generations (van der Graaf et al., 2015, Shahryary et al., 2020). This raises the possibility that these selected sites may contribute to long-term adaptation in a classical Darwinian framework, an intriguing hypothesis that warrants further investigation.

Collectively, this study improves our basic understanding of how environmental signals shaped DNA methylome dynamics in Arabidopsis. Future research should extend our analysis to functional validation through targeted methylome editing techniques, explore multigenerational inheritance explicitly, and evaluate translational potential in crop breeding programs aimed at improving stress resilience.

## Data availability statement

The data that support the findings of this study are available in article supplementary material and on request from corresponding authors.

## Author Contributions

Experiments were designed by FJ, analysis were performed by RB, JJGF, RRH, research articles were searched by FW and RRH, data interpretation was performed by RB, CL, FJ, paper written by RB with input from CL, FJ and JPS. Epimutation analysis was performed by ZZ. Written paper was reviewed by all the co-authors.

## Funding statement

This work was supported (RB & CL) by the EpiCrossBorders: International Hemholtz-Edinburgh Research School for Epigenetics, which promotes cross-border research in epigenetics.

## Conflict of interest disclosure

not declared

## Notes

### Competing Interest Statement

The authors have declared no competing interest.

