## Supporting Information for "DNA methylome responses to biotic and abiotic stress in *Arabidopsis thaliana*: A multi-study analysis"

Affiliations here
[1]Research Unit Environmental Simulation (EUS), Helmholtz Munich, Neuherberg, Germany
[2]International Center for Tropical Agriculture (CIAT), CGIAR, Cali, Colombia
[3]Plant Epigenomics, TUM School of Life Sciences, Technical University of Munich, Freising, Germany
[4]Institute of Lung Health and Immunity (LHI), Comprehensive Pneumology Center (CPC-M), Helmholtz Munich, Neuherberg, Germany

[#]These authors contributed equally to this work
*Corresponding authors

**Supplementary Figures**

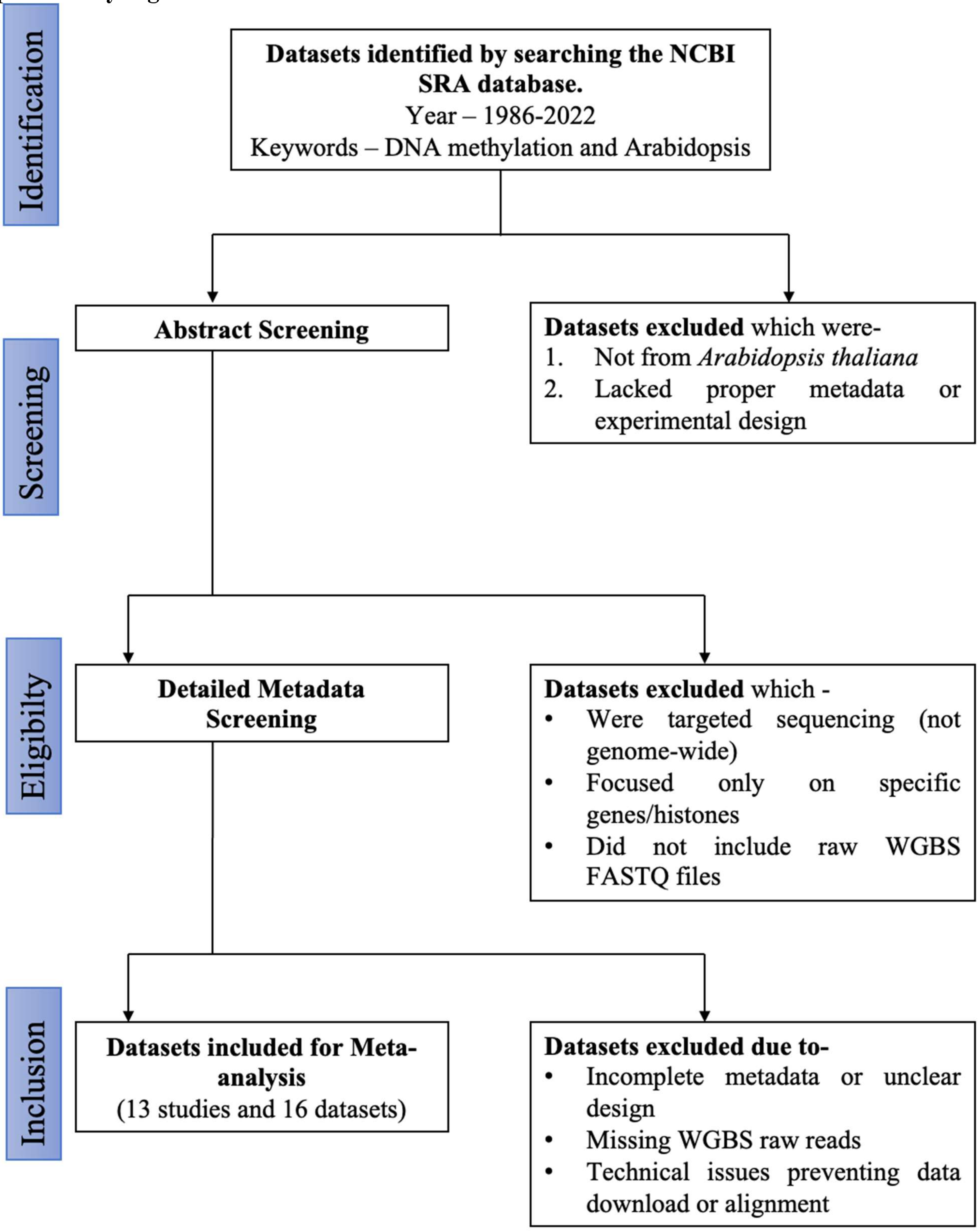


**Supplementary Figure S1** – Schematic representation of dataset selection and filtering process for *Arabidopsis thaliana* WGBS experiments identified from the NCBI SRA database and included in the meta-analysis.

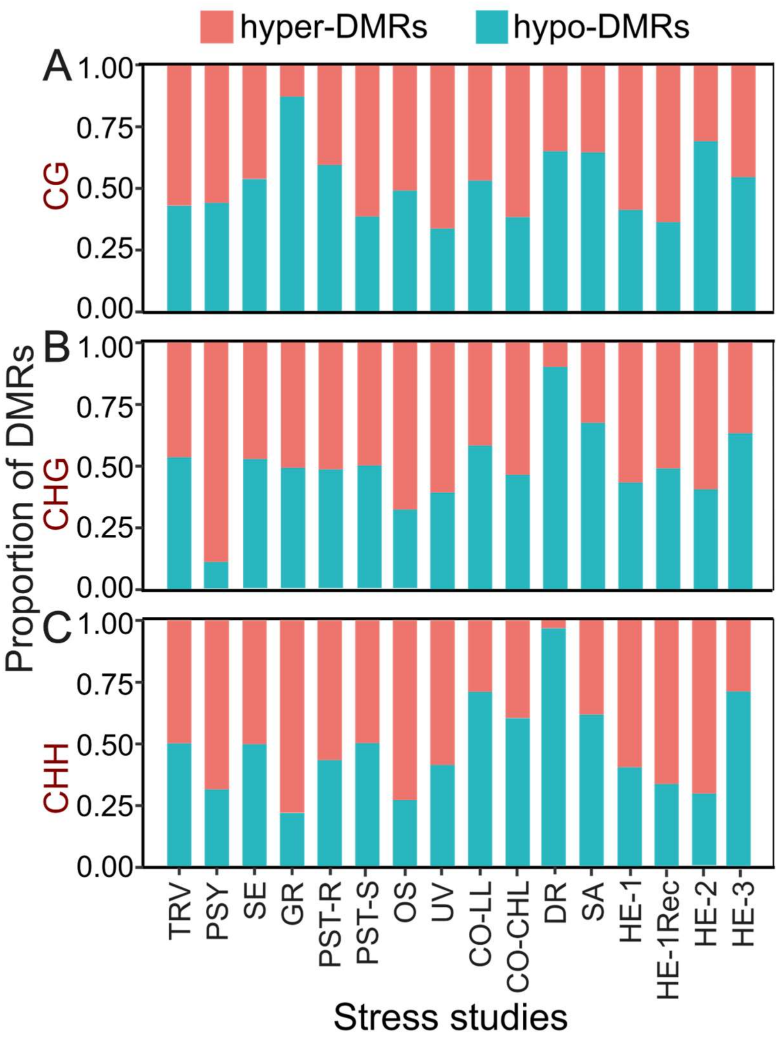


**Supplementary Figure S2** – Proportions of hyper- and hypo-methylated regions in stress-related DMRs. Each bar represents the relative proportions of hyper- and hypo-methylated regions for specific study for **(A)** CG, **(B)** CHG and **(C)** CHH contexts.

**A**

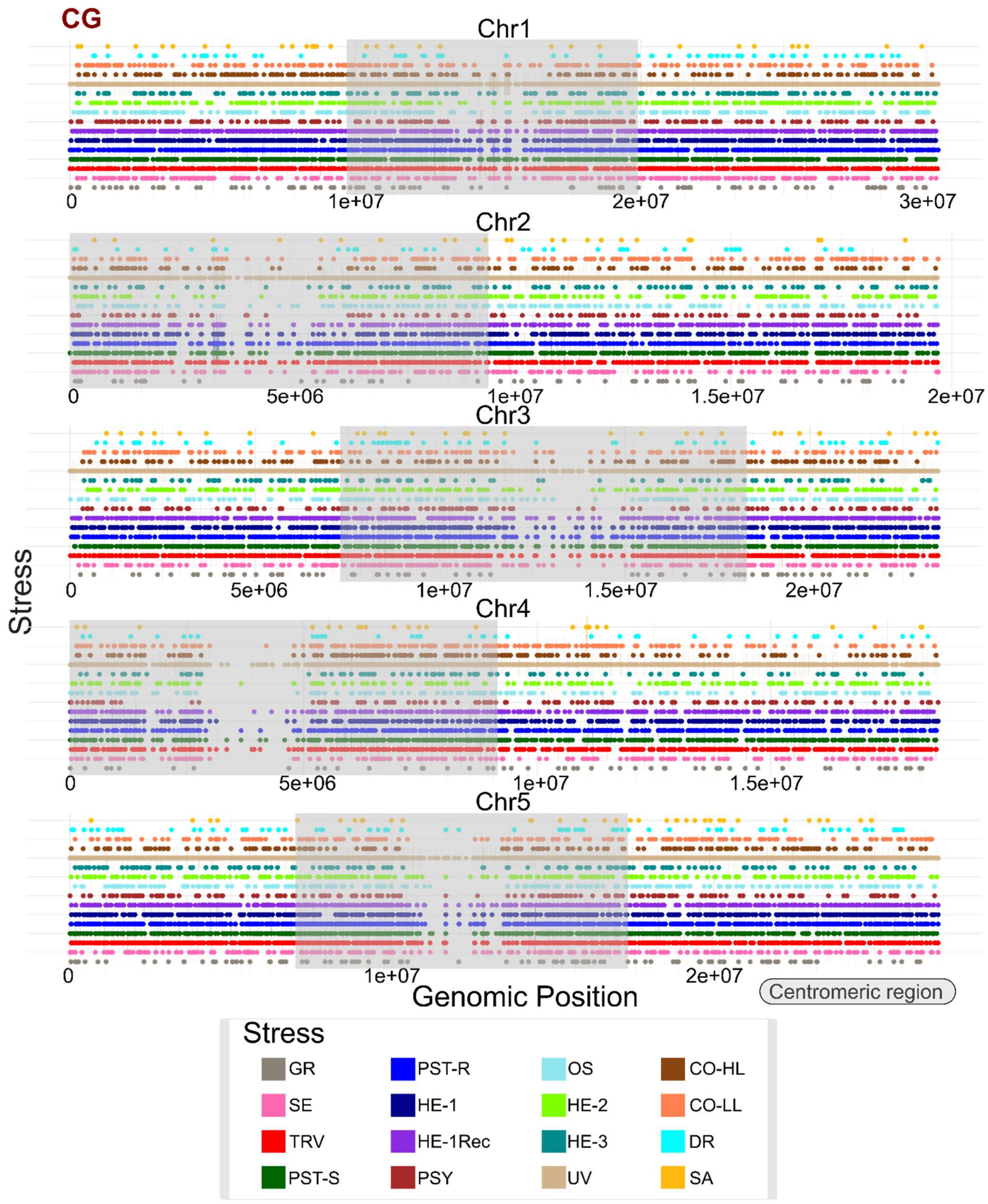
CG
Chr1
0
1e+07
2e+07
3e+07
Chr2
0
5e+06
1e+07
1.5e+07
2e+07
Chr3
0
5e+06
1e+07
1.5e+07
2e+07
Chr4
0
5e+06
1e+07
1.5e+07
Chr5
0
1e+07
2e+07
Stress
Genomic Position
Centromeric region
Stress
GR
PST-R
OS
CO-HL
SE
HE-1
HE-2
CO-LL
TRV
HE-1Rec
HE-3
DR
PST-S
PSY
UV
SA

**B**

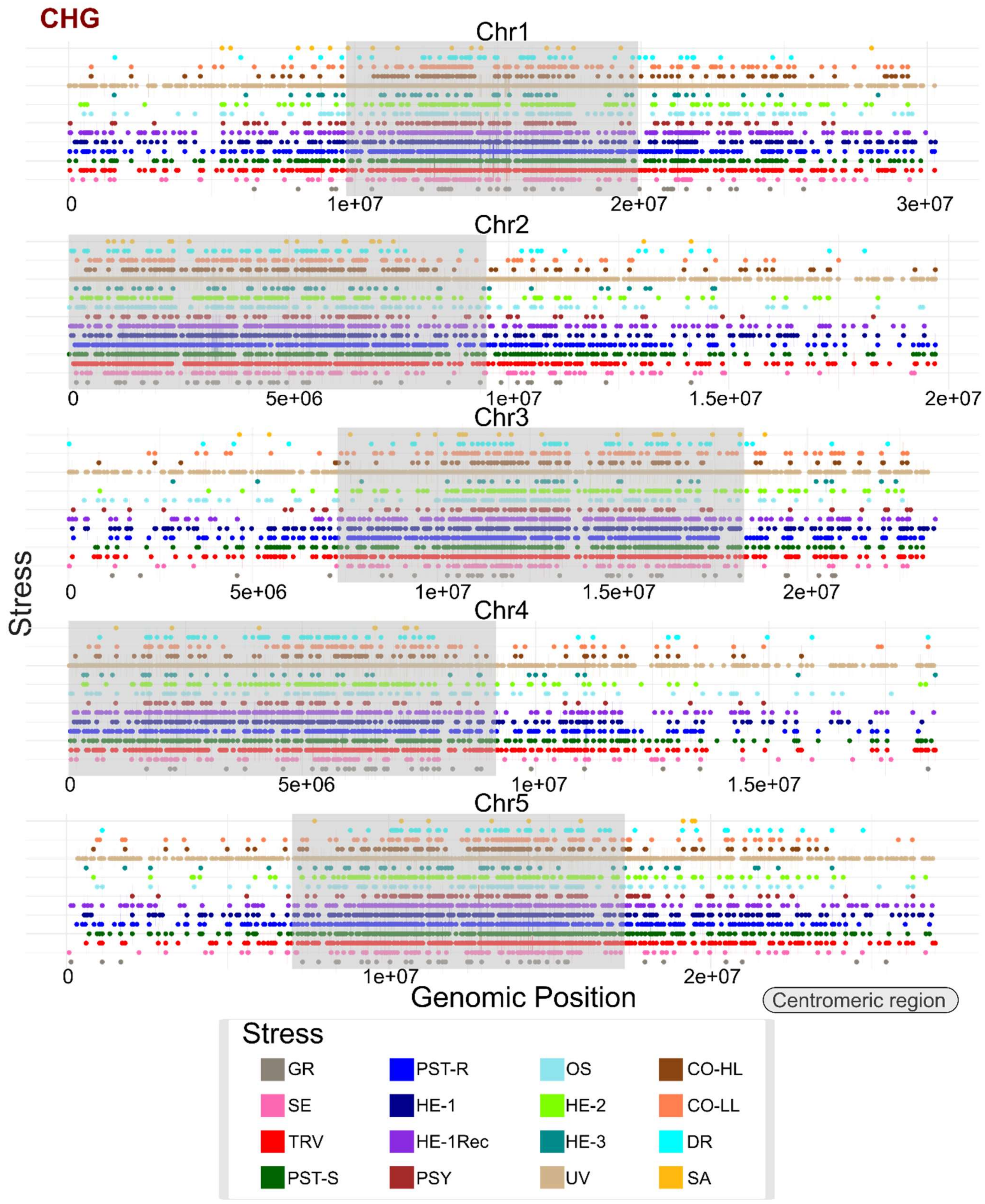
CHG
Chr1
0
1e+07
2e+07
3e+07
Chr2
0
5e+06
1e+07
1.5e+07
2e+07
Chr3
0
5e+06
1e+07
1.5e+07
2e+07
Chr4
0
5e+06
1e+07
1.5e+07
Chr5
0
1e+07
2e+07
Stress
Genomic Position
Centromeric region
Stress
GR
PST-R
OS
CO-HL
SE
HE-1
HE-2
CO-LL
TRV
HE-1Rec
HE-3
DR
PST-S
PSY
UV
SA

**C**

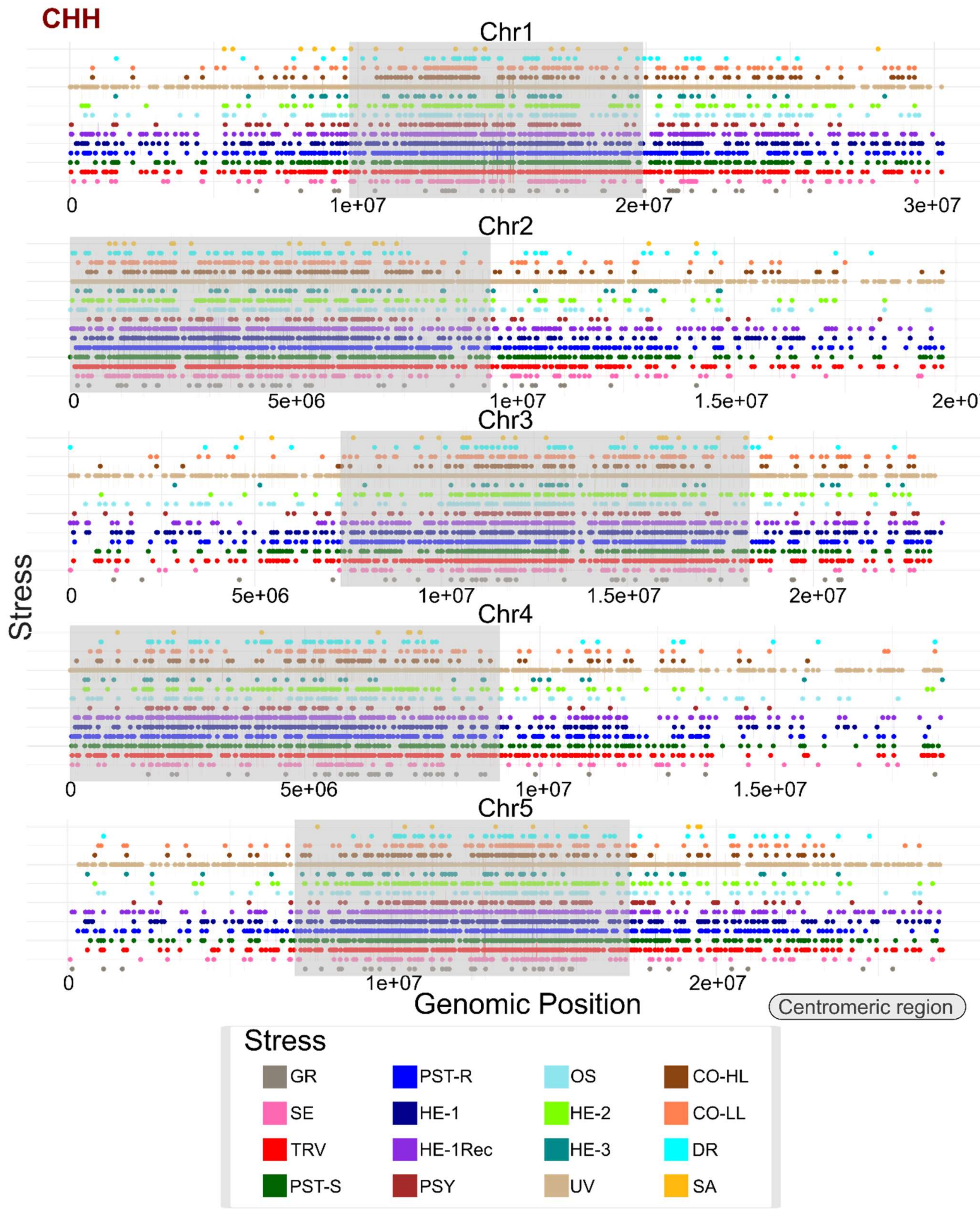


**Supplementary Figure S3 - Genome-wide distribution of stress-responsive DMRs across methylation contexts.** (A) **CG**, (B) **CHG**, and (C) **CHH**. Within each panel, the five horizontal tracks correspond to chromosomes 1–5 (top to bottom). Each point marks the genomic position (midpoint) of a DMR detected in a given stress study; colours indicate the stress/study as shown in the key. Grey shaded boxes highlight the annotated centromeric/pericentromeric regions for each chromosome.

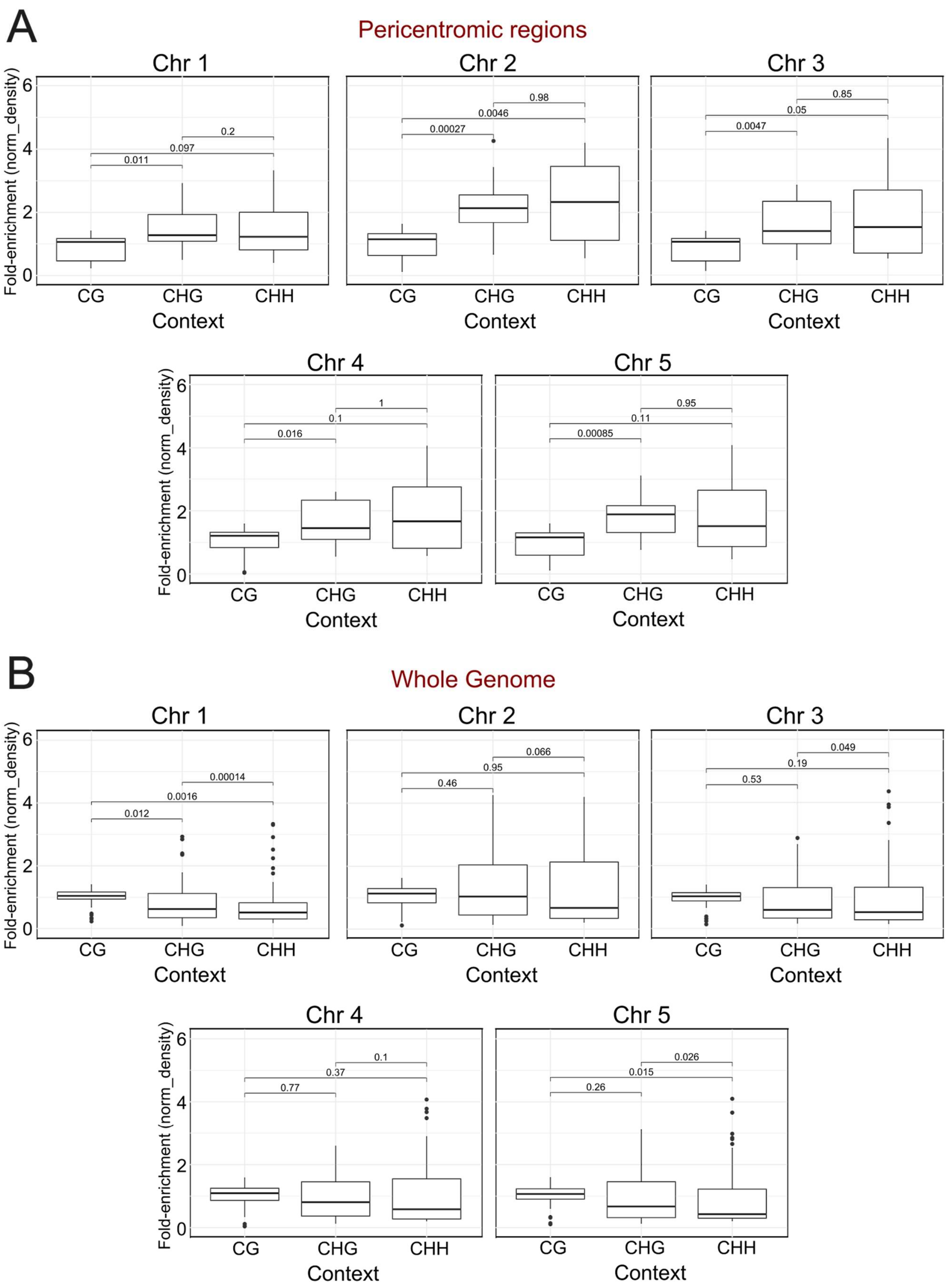
A
Pericentromic regions
Chr 1
Chr 2
Chr 3
Chr 4
Chr 5
Fold-enrichment (norm_density)
Context
CG
CHG
CHH
0.011
0.097
0.2
0.00027
0.0046
0.98
0.0047
0.05
0.85
0.016
0.1
1
0.00085
0.11
0.95
B
Whole Genome
Chr 1
Chr 2
Chr 3
Chr 4
Chr 5
Fold-enrichment (norm_density)
Context
CG
CHG
CHH
0.012
0.0016
0.00014
0.46
0.95
0.066
0.53
0.19
0.049
0.77
0.37
0.1
0.26
0.015
0.026

**Supplementary Figure S4 - Context-specific DMR density in pericentromeric regions versus genome-wide.**

Boxplots summarize the distribution of **normalized DMR density** (fold-enrichment) for **CG**, **CHG**, and **CHH** contexts. The genome was tiled into **non-overlapping 500-kb windows** (Chr1–Chr5). For each window, DMR density was computed as the number of **significant DMRs** overlapping the window divided by window size, and then **normalized by the genome-wide mean** of that context (yielding a fold-change; 1 = genome average). **(A) Pericentromeric region:** windows within the pericentromere of each chromosome are shown; **(B) Whole genome:** all windows on the chromosome are shown. Boxes show medians and interquartile ranges across windows; whiskers extend to 1.5× IQR, with points indicating outliers. **Paired Wilcoxon signed-rank tests** (across matched windows) compare contexts; formatted *p*-values are displayed above comparisons.

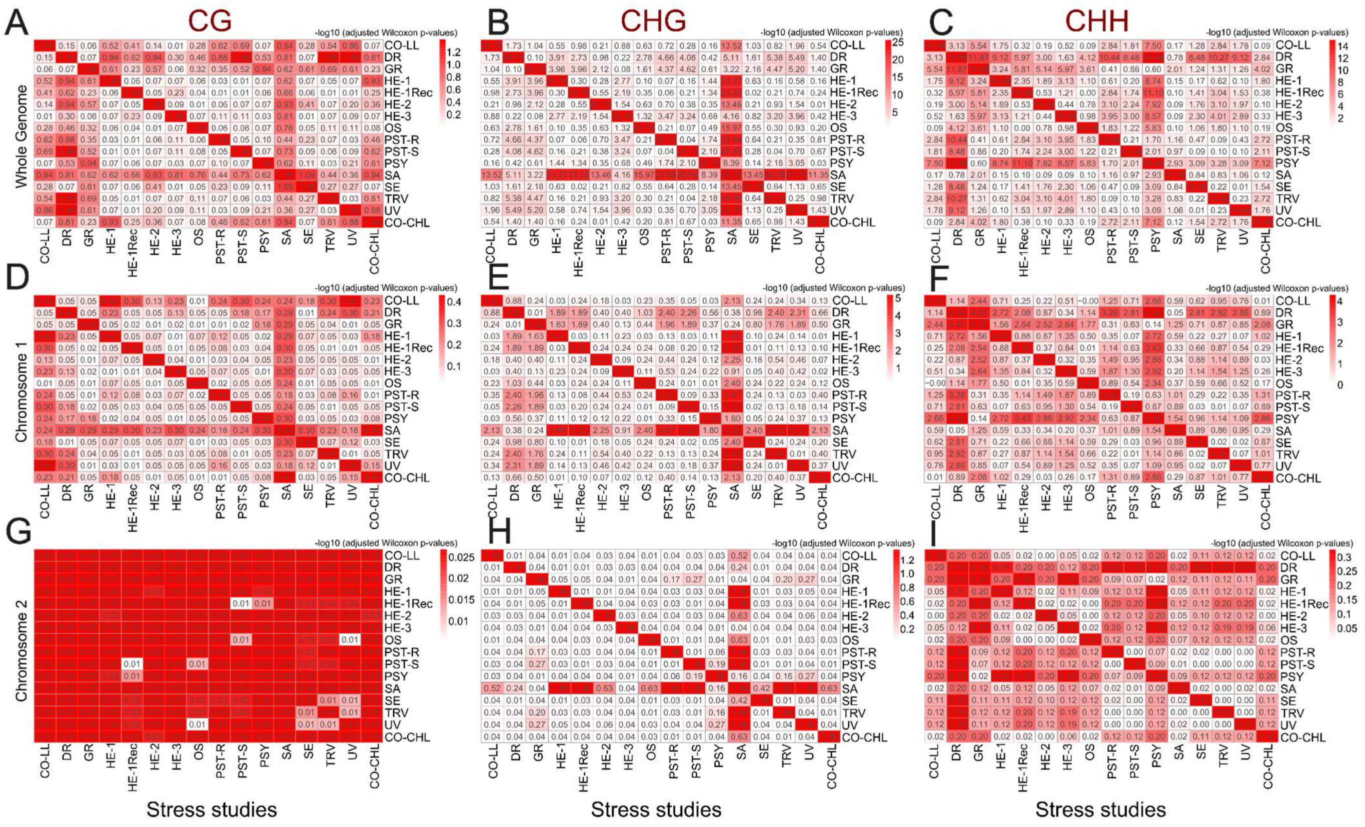

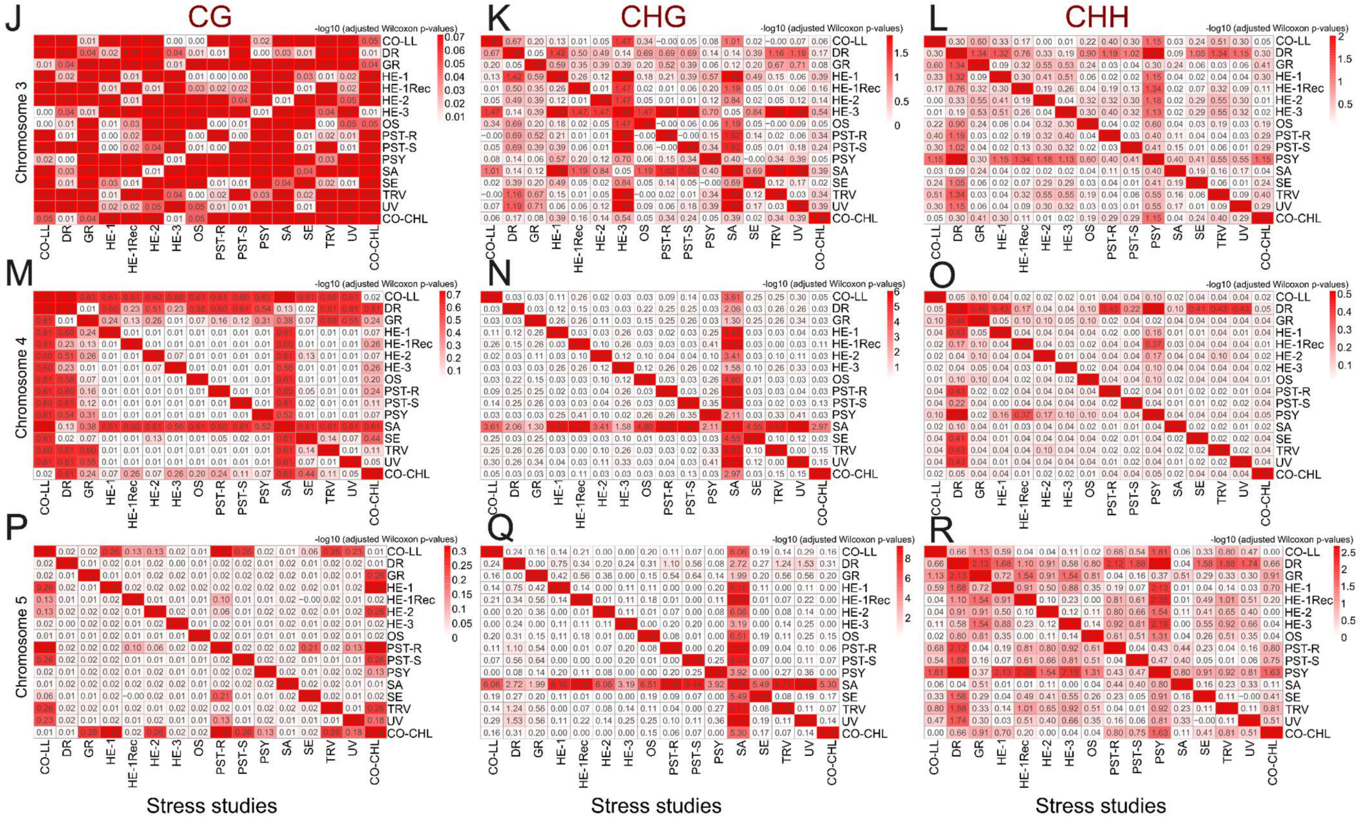

**Supplementary Figure S5 - Pairwise differences in DMR density across stress studies (Kruskal–Wallis framework).** Heatmaps show pairwise comparisons of DMR density between studies for each methylation context—**CG** (left column), **CHG** (middle), and **CHH** (right)—separately by chromosome: **Chr1** (D-F), **Chr2** (G-I), **Chr3** (J-L), **Chr4** (M-O), **Chr5** (P-R), and **Genome-wide (all chromosomes pooled)** (A-C). For each chromosome/context, DMR density was tested across studies with a Kruskal–Wallis rank-sum test; when the overall test was significant, post-hoc pairwise Wilcoxon with Benjamini–Hochberg correction. Matrix cells display the adjusted *p*-values (numbers in cells), and colour encodes −log10 (adjusted *p*) (white = not significant/high *p*; dark red = stronger evidence of difference/low *p*). Diagonals are self-comparisons. Study abbreviations follow the legend used in the main figures.

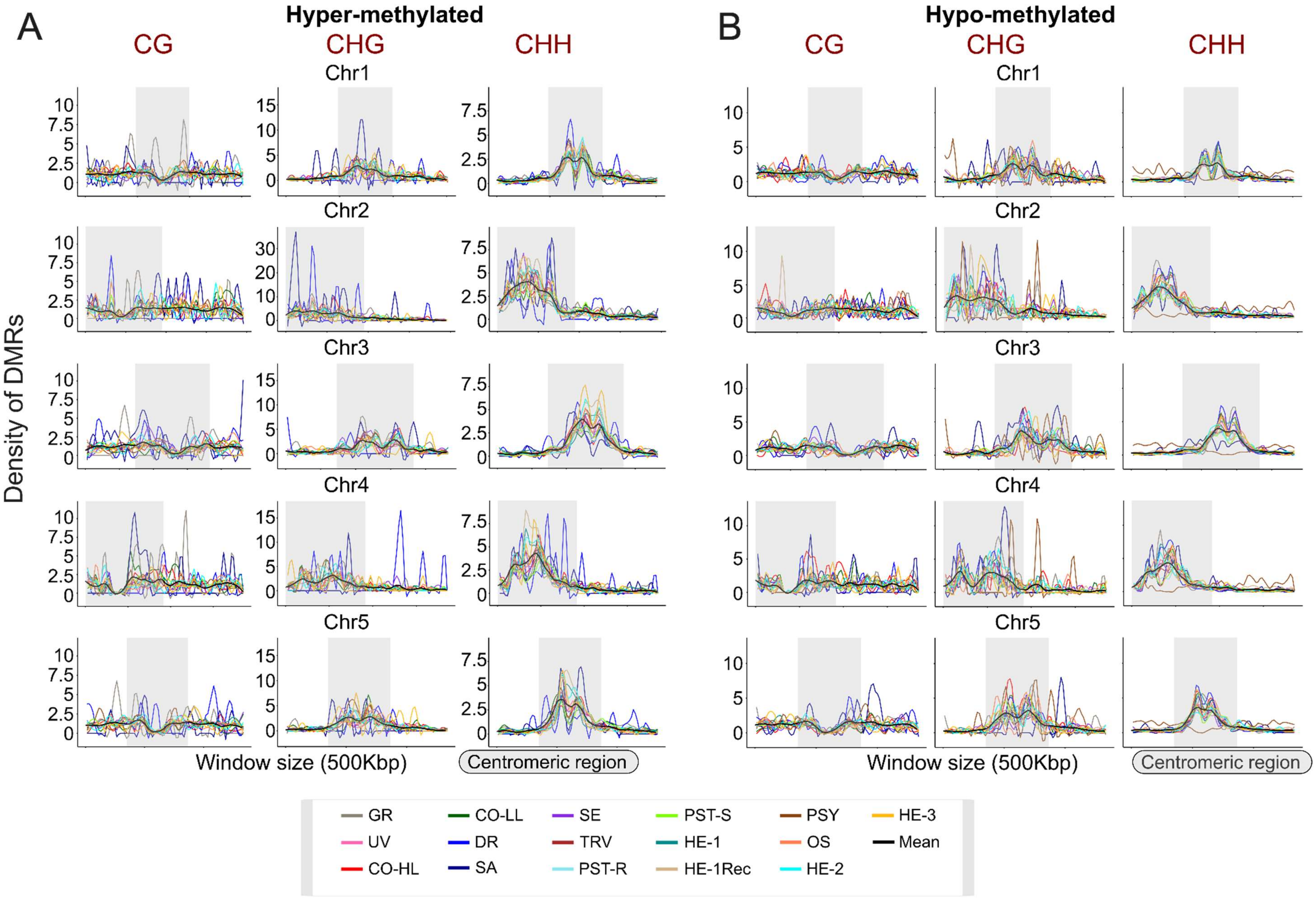
A
Hyper-methylated
CG
CHG
CHH
B
Hypo-methylated
CG
CHG
CHH
Chr1
Chr2
Chr3
Chr4
Chr5
Density of DMRs
Window size (500Kbp)
Centromeric region
Window size (500Kbp)
Centromeric region
GR
UV
CO-HL
CO-LL
DR
SA
SE
TRV
PST-R
PST-S
HE-1
HE-1Rec
PSY
OS
HE-2
HE-3
Mean

**Supplementary Figure S6-** Density of DMRs per genomic window across studies. Densities are shown for three methylation contexts: CG, CHG, and CHH. The gray segment highlights the centromeric region, where DMR density tends to peak in most contexts. Each line represents a different study, as indicated by the legend. **(A)** Hyper-methylated **(B)** Hypo-methylated.

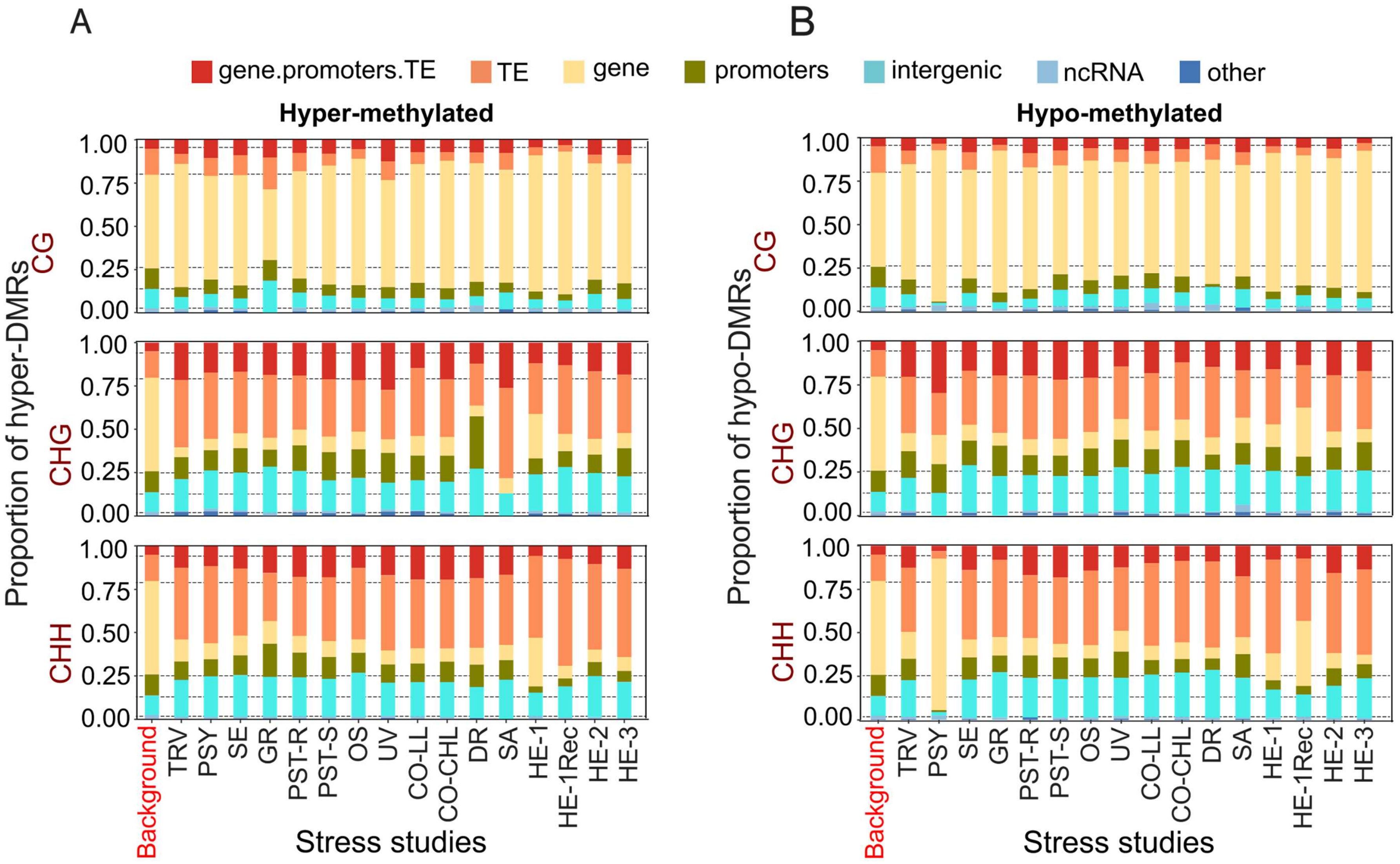

A
B
gene.promoters.TE
TE
gene
promoters
intergenic
ncRNA
other
Hyper-methylated
Hypo-methylated
Proportion of hyper-DMRs
Proportion of hypo-DMRs
CG
CHG
CHH
1.00
0.75
0.50
0.25
0.00
Background
TRV
PSY
SE
GR
PST-R
PST-S
OS
UV
CO-LL
CO-CHL
DR
SA
HE-1
HE-1Rec
HE-2
HE-3
Stress studies

**Supplementary Figure S7-** Proportion of DMRs annotated to different genomic features (e.g., gene body, promoter, intergenic regions, etc.) for each study. The three methylation contexts (CG, CHG, CHH) are analyzed separately. The "Background" bar represents the genomic distribution of features across the entire genome for comparison. **(A)** Hyper-methylated **(B)** Hypo-methylated.

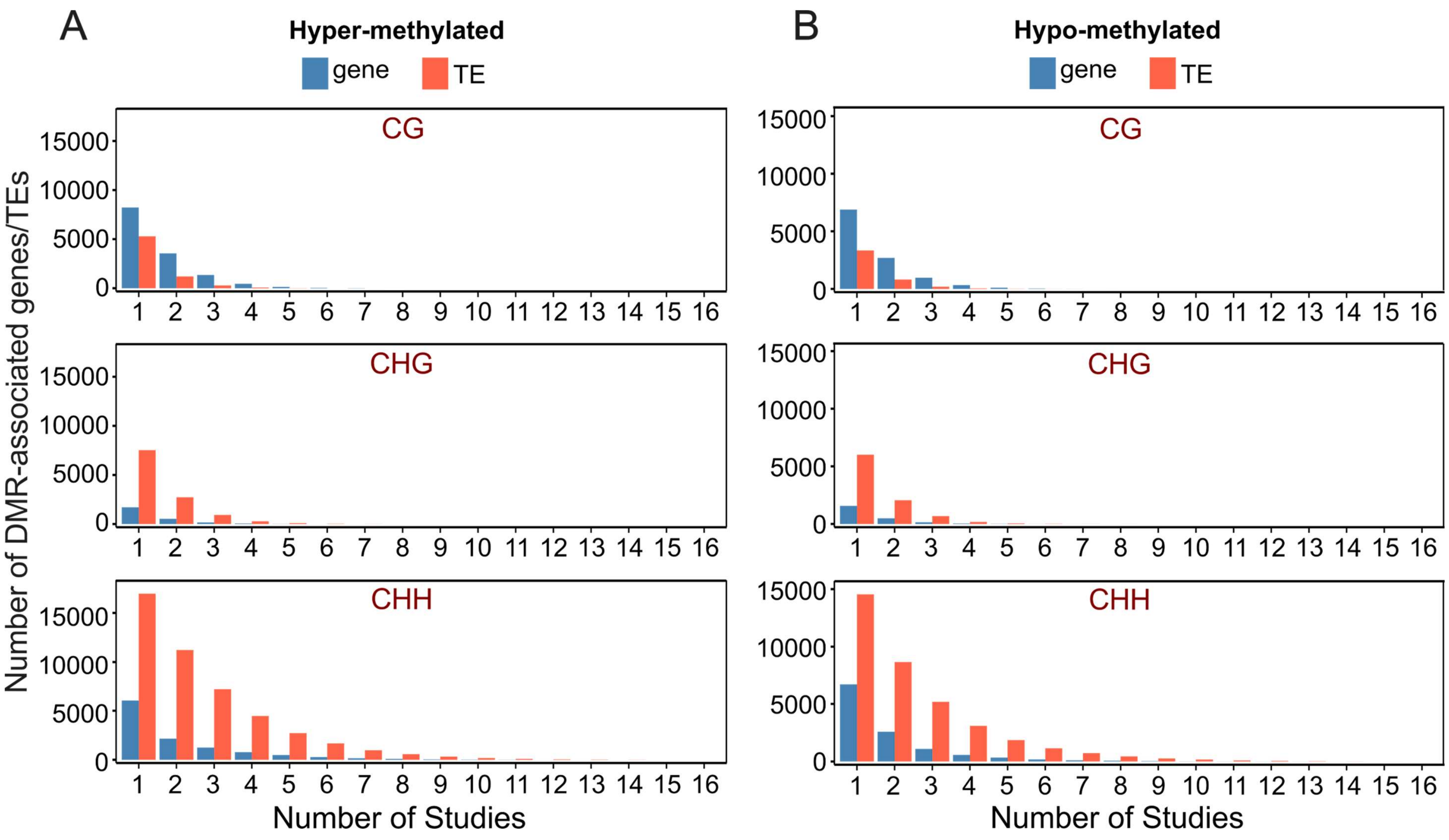

**Supplementary Figure S8**- Number of DMRs (hyper and hypo) -associated genes and TEs shared across studies. Bar plots show the number of DMR-associated genes (blue) and TEs (red) identified in pairwise comparisons of studies. Each panel corresponds to one of the three methylation contexts: CG, CHG, and CHH. The x-axis represents the number of studies in which DMR-associated features are shared, while the y-axis indicates the count of shared genes or TEs. **(A)** Hyper-methylated **(B)** Hypo-methylated.

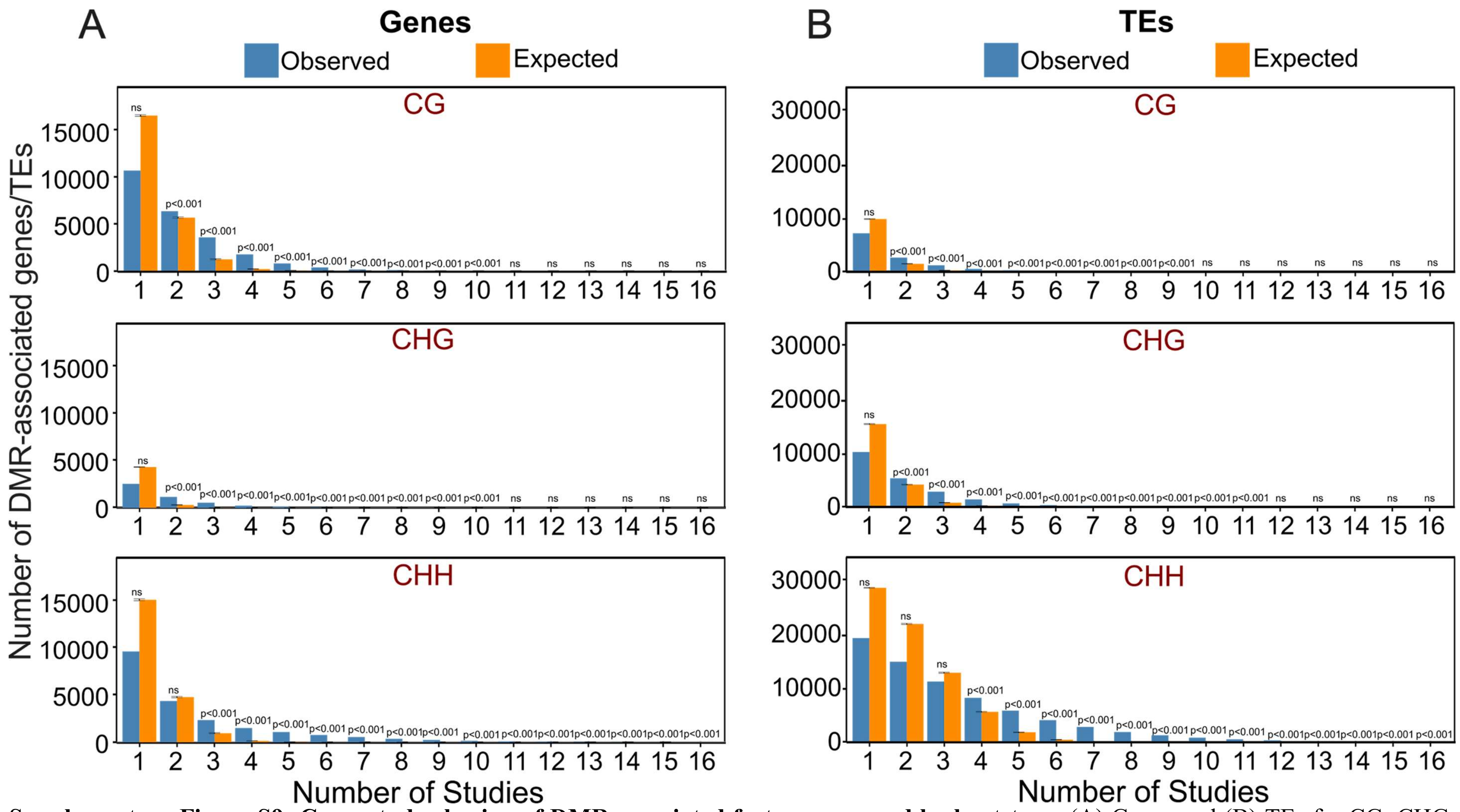


**Supplementary Figure S9- Cross-study sharing of DMR-associated features assessed by bootstrap.** (A) Genes and (B) TEs for CG, CHG, and CHH. Blue bars show the observed number of unique features present in 16 studies (x-axis). Orange bars summarize the expected counts under a bootstrap null that preserves each study's list size (10,000 resamples drawn from the background set); black error bars indicate the 95% bootstrap interval. Text above bars gives one-sided enrichment *p*-values computed as the fraction of bootstrap replicates with counts ≥ the observed ("n.s.", not significant).

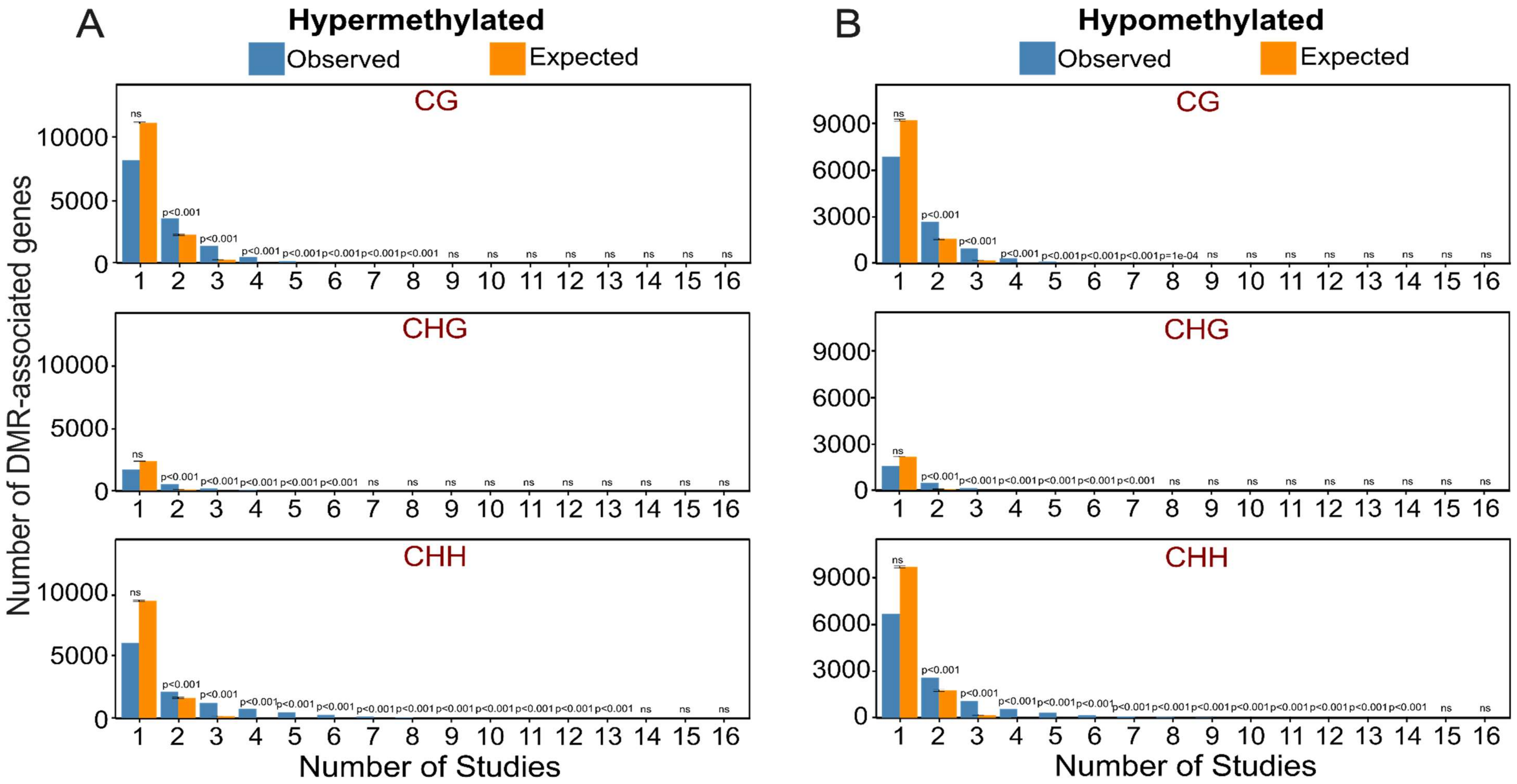


**Supplementary Figure S10 - Cross-study sharing of DMR-associated features assessed by bootstrap.** (A) Hypermethylated and (B) Hypomethylated genes for CG, CHG, and CHH. Blue bars show the observed number of unique features present in 16 studies (x-axis). Orange bars summarize the expected counts under a bootstrap null that preserves each study's list size (10,000 resamples drawn from the background set); black error bars indicate the 95% bootstrap interval. Text above bars gives one-sided enrichment *p*-values computed as the fraction of bootstrap replicates with counts ≥ the observed ("n.s.", not significant).

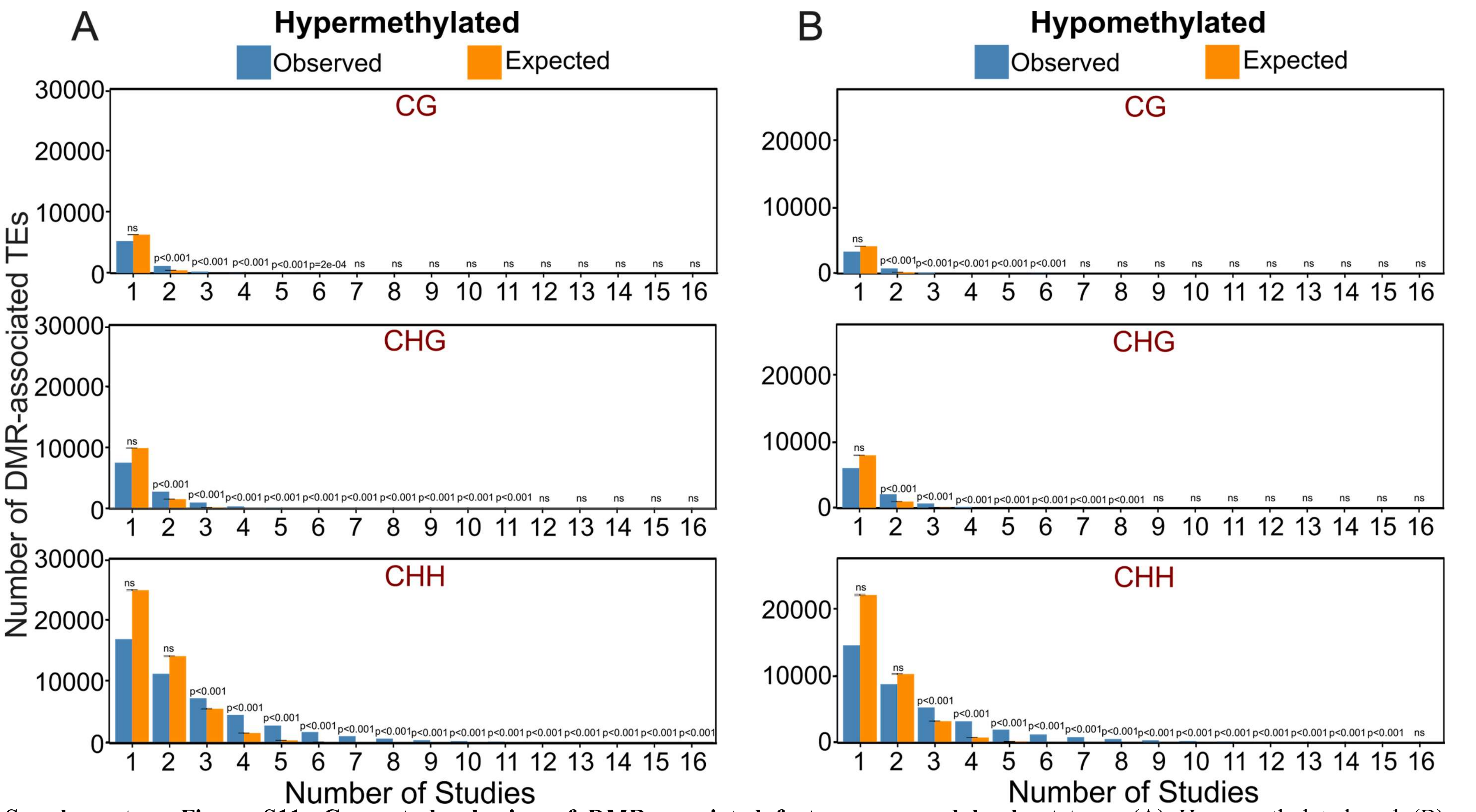


**Supplementary Figure S11- Cross-study sharing of DMR-associated features assessed by bootstrap.** (A) Hypermethylated and (B) Hypomethylated TEs for CG, CHG, and CHH. Blue bars show the observed number of unique features present in 16 studies (x-axis). Orange bars summarize the expected counts under a bootstrap null that preserves each study's list size (10,000 resamples drawn from the background set); black error bars indicate the 95% bootstrap interval. Text above bars gives one-sided enrichment *p*-values computed as the fraction of bootstrap replicates with counts ≥ the observed ("n.s.", not significant).

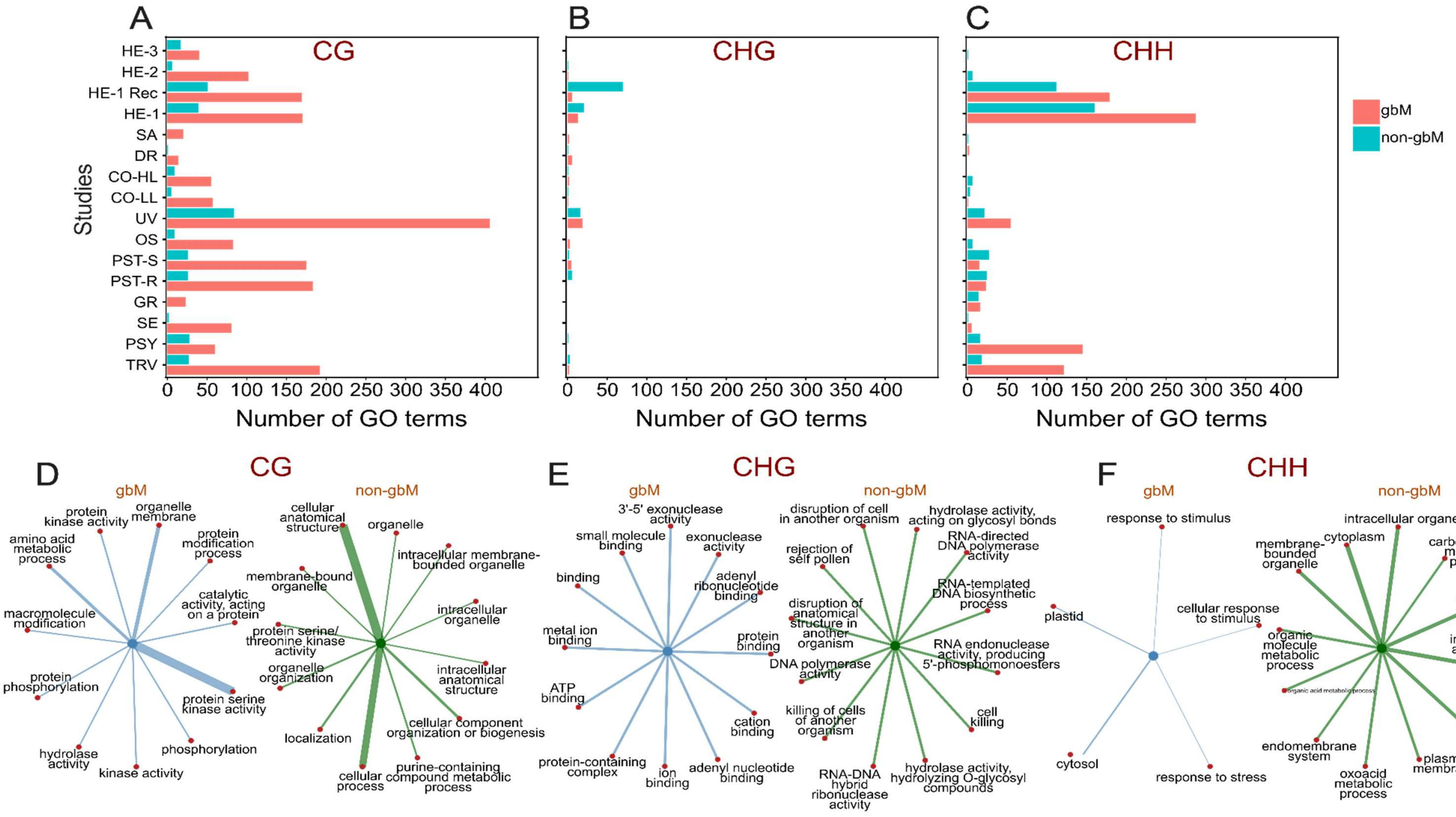


**Supplementary Figure S12 - gbM vs non-gbM composition of DMR-associated genes and their GO enrichments. (A–C)** For each methylation context (CG, CHG, CHH; panel headers), horizontal bars show, per study (y-axis), the number of DMR-associated genes classified as gbM vs non-gbM (colours as in the inset key). **(D–F)** Networks of enriched GO terms for the same contexts, shown separately for gbM (left, blue) and non-gbM (right, green). The central node denotes the gene class; red nodes are GO terms. Edge thickness is proportional to the odds ratio from Fisher's exact test; thicker edges indicate stronger enrichment. Unless otherwise noted, only the top 12 terms by odds ratio are displayed for readability, and terms meeting FDR < 0.05 (Benjamini–Hochberg) are considered significant. *Note: No term passed FDR in CHG context, the top 12 terms by odds ratio are shown for visualization only (no significance implied).*

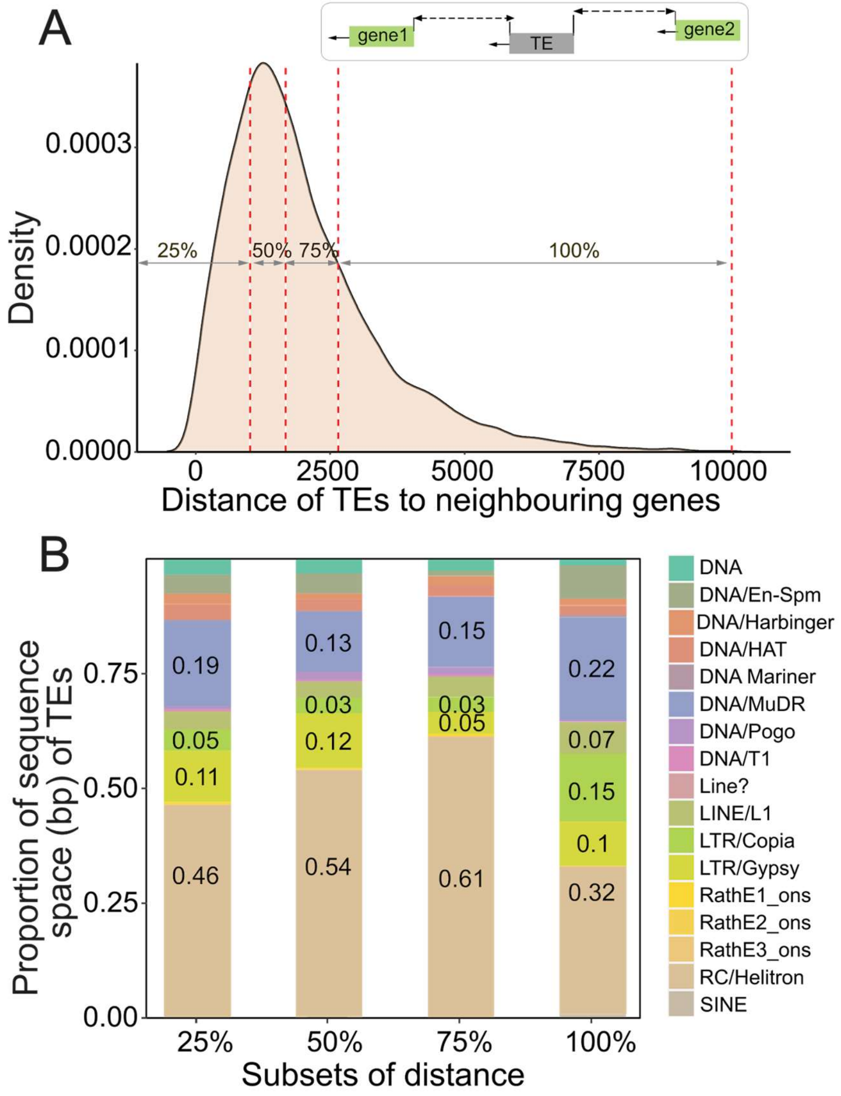


**Supplementary Figure S13 - (A)** Density plot showing the distribution of TEs in proximity to genes. The x-axis represents the relative distance of TEs from nearby genes, with 0% and 100% corresponding to the start and end of the gene regions, respectively. Red dashed lines indicate quartiles (25%, 50%, and 75%). **(B)** Stacked bar chart representing the enrichment of TE superfamilies across distance subsets of DMR-associated genes. The y-axis shows the proportion of the sequence space (in base pairs, bp) occupied by each TE superfamily.

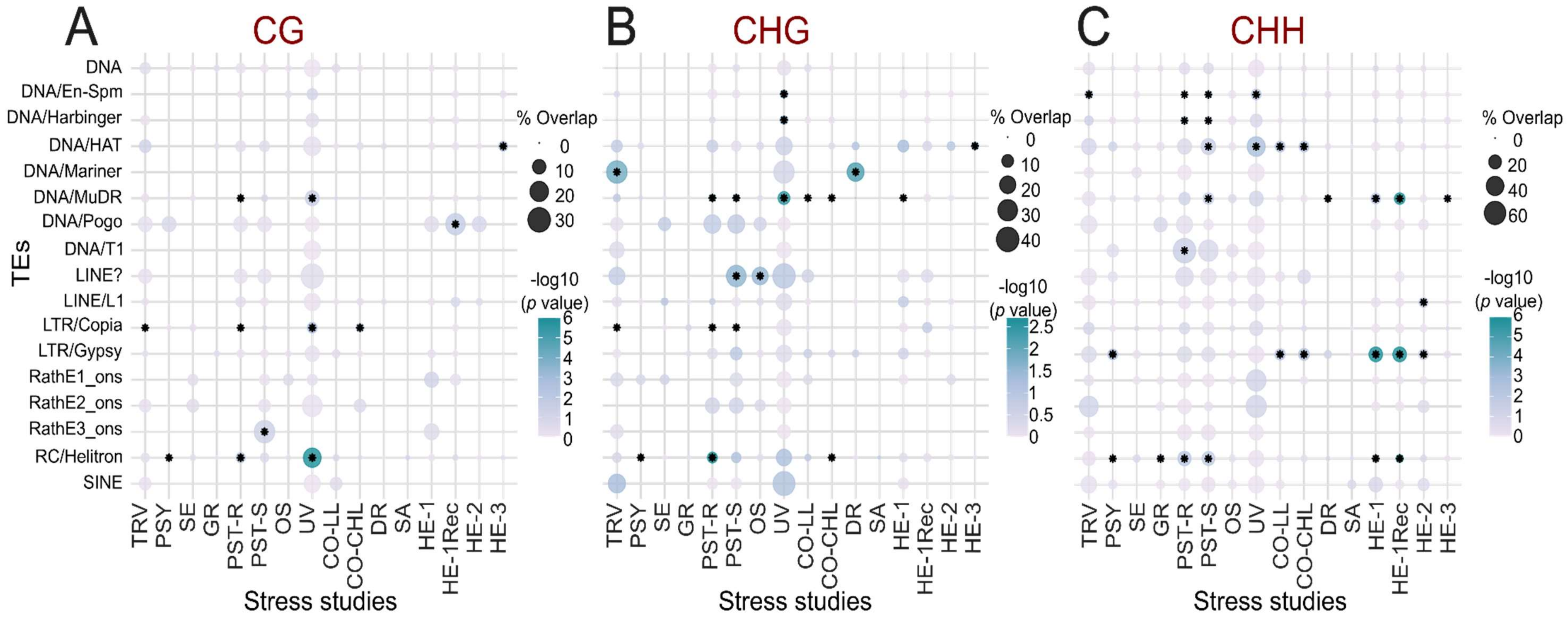


**Supplementary Figure S14 – Permutation analysis of gene-proximal TE superfamilies with context-specific DMRs across stresses. (A)** CG, **(B)** CHG, **(C)** CHH. Columns are stress studies and rows are TAIR10 TE superfamilies. Each circle summarizes, for a given study and superfamily, the percentage of gene-proximal TEs (elements within ±1.2 kb of gene TSSs) that overlap at least one DMR in that context; bubble area is proportional to this percentage. Colour encodes −log10 of the two-sided permutation *p*-value obtained by shuffling DMR labels within the proximal-TE universe (10,000 permutations per study); black stars mark $p < 0.05$. The TE universe was derived from TAIR10 annotations and then restricted to gene-proximal elements.

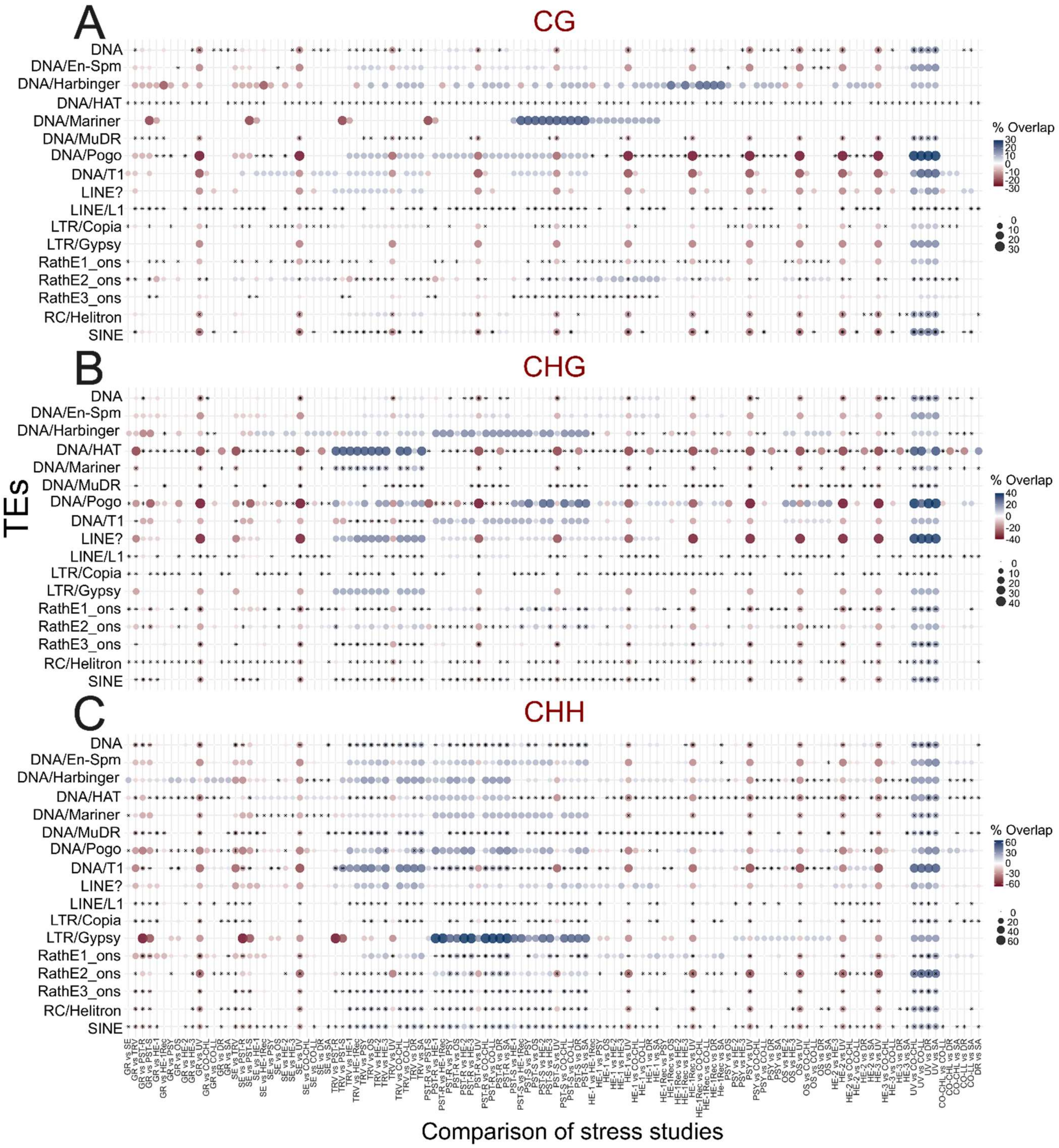


**Supplementary Figure S15: Pairwise differences in enrichment of gene-proximal TE superfamilies with DMRs across studies. (A)** CG, **(B)** CHG, **(C)** CHH. Rows are TAIR10 TE superfamilies; columns are all pairwise study comparisons. For each superfamily and comparison, the tile shows % of gene-proximal TEs (Δ%\Delta\%Δ%) in that superfamily that overlap ≥1 DMR in study a) − (**% in study b). Colour encodes the signed Δ%\Delta\%Δ% - diverging scale; red = a > b, blue = a < b; white ≈ 0). Stars mark two-sided permutation significance ($p < 0.05$) obtained by shuffling study labels within the pooled proximal-TE set 10,000 times and recomputing Δ%\Delta\%Δ%. Gene-proximal TEs were defined as elements within ±1.2 kb of genes.

**Supplementary Files**

**Supplementary File S1 — Shared gene/TE lists across studies and methylation contexts**
For each methylation context (CG, CHG, CHH) and feature type (Gene or TE), the table summarizes elements shared by multiple studies.
Columns:

- Shared_By – Category describing how many studies share the feature (e.g., "All genes", "2 studies", …).
- Count – Number of genes/TEs in that category.
- Feature – "Gene" or "TE".
- Context – Methylation context (CG/CHG/CHH).
- Names – Comma-separated identifiers (AGI for genes; TAIR/TE IDs for TEs).
- Function – Placeholder for functional notes (empty in the provided file).

**Supplementary File S2 — CG context: GO term frequency summary**
Columns:

- term_name – GO term name.
- Exact_Count – Number of study-specific enrichment lists in which the term appears (CG context).
- Cumulative_Count – Running/rank order used for plotting summaries.

**Supplementary File S3 — CHG context: GO term frequency summary**
Columns: term_name; Exact_Count; Cumulative_Count (as in S2, for CHG).
**Supplementary File S4 — CHH context: GO term frequency summary**
Columns: term_name; Exact_Count; Cumulative_Count (as in S2, for CHH).

**Supplementary File S5 — GO term sharing networks between studies (edge lists)**
Edge lists used to build study-to-study GO-sharing networks for each methylation context (CG, CHG, CHH). Organized in three 5-column blocks (one per context).
Per-context columns:

- from – Source study.
- to – Target study.
- weight – Number of GO terms shared between the two studies in that context.
- study – Study for which go_count is reported on that row.
- go_count – Total GO term count in the enrichment list for the study named in study.

**Supplementary File S6 — CG functional categories**
Grouping of CG-context GO terms into broader functional classes used for visualization (e.g., "Binding properties," "Enzymatic activity," "Epigenetic mechanisms," "Transport function," "Reproduction and Development"). An Excluded flag marks classes/terms removed before plotting to avoid redundancy.
Columns:

- CG context – Curated high-level functional class name.
- Additional columns – Reserved for flags/notes (e.g., Excluded).

**Supplementary File S7 — CHG functional curation**
As in S6 but for the CHG context, listing high-level classes assigned to CHH GO terms; an Excluded marker indicates terms not retained for visualization.

**Supplementary File S8 — CHH functional curation**
As in S6 but for the CHH context, listing high-level classes assigned to CHH GO terms; an Excluded marker indicates terms not retained for visualization.

**Supplementary File S9 — Hyper-DMR shared gene lists across studies**
For each methylation context (CG/CHG/CHH), lists genes present in hyper-methylated DMR-associated gene sets and indicates how many studies share each category.
Columns:

- Hyper-DMRs – sharing category (e.g., "All genes", "2 studies", …)
- Count – number of genes in that category
- Feature – feature type ("Gene"and "TE")
- Context – methylation context (CG/CHG/CHH)
- Names – comma-separated AGI identifiers

**Supplementary File S10 — Hypo-DMR shared gene lists across studies**
Same structure as S9 but for hypo-methylated DMR-associated gene sets.
Columns: Hypo-DMRs, Count, Feature, Context, Names (AGIs).

**Supplementary File S11 — Master index of genes and TEs across studies and contexts**
Consolidated list of every ID (gene AGI or TE ID) detected in the DMR annotation-overlap analysis across all studies and contexts.
Columns:

- ID – gene (AGI) or TE identifier
- Count – in how many input files the ID appears (1–16)
- Feature – "Gene" or "TE"
- Context – methylation context (CG/CHG/CHH)

**Supplementary File S12 — Figure 8 gene sets with linked TEs (by context)**
Worksheets:

- CG_genes_Figure8, CHG_genes_Figure8, CHH_genes_Figure8 — per-context tables of genes featured in Figure 8, with the number of studies and linked TE(s).
  Columns: gene_id (AGI), studies_seen, TE_ids, TE_superfamilies, studies (study abbreviations).
- CG-genelist-figure8, CHG-genelist-figure8, CHH-genelist-figure8 — mapping tables from AGI to display name/description and identifier namespaces.
  Columns: initial_alias, converted_alias, name, description, namespace.
